# Exploring masses and internal mass distributions of single carboxysomes in free solution using fluorescence and interferometric scattering in an anti-Brownian trap

**DOI:** 10.1101/2022.08.23.505029

**Authors:** Abhijit A. Lavania, William B. Carpenter, Luke M. Oltrogge, Davis Perez, Julia S. Borden, David F. Savage, W. E. Moerner

## Abstract

Carboxysomes are self-assembled bacterial microcompartments that facilitate carbon assimilation by co-localizing the enzymes of CO_2_ fixation within a protein shell. These microcompartments can be highly heterogeneous in their composition and filling, so measuring the mass and loading of an individual carboxysome would allow for better characterization of its assembly and function. To enable detailed and extended characterizations of single nanoparticles in solution, we recently demonstrated an improved Interferometric Scattering Anti-Brownian ELectrokinetic (ISABEL) trap, which tracks the position of a single nanoparticle via its scattering of a near-infrared beam and applies feedback to counteract its Brownian motion. Importantly, the scattering signal can be related to the mass of nanoscale proteinaceous objects, whose refractive indices are well-characterized. We calibrate single-particle scattering cross-section measurements in the ISABEL trap and determine individual carboxysome masses in the 50-400 MDa range by analyzing their scattering cross-sections with a core-shell model. We further investigate carboxysome loading by combining mass measurements with simultaneous fluorescence reporting from labeled internal components. This method may be extended to other biological objects, such as viruses or extracellular vesicles, and can be combined with orthogonal fluorescence reporters to achieve precise physical and chemical characterization of individual nanoscale biological objects.

**TOC Graphic:** 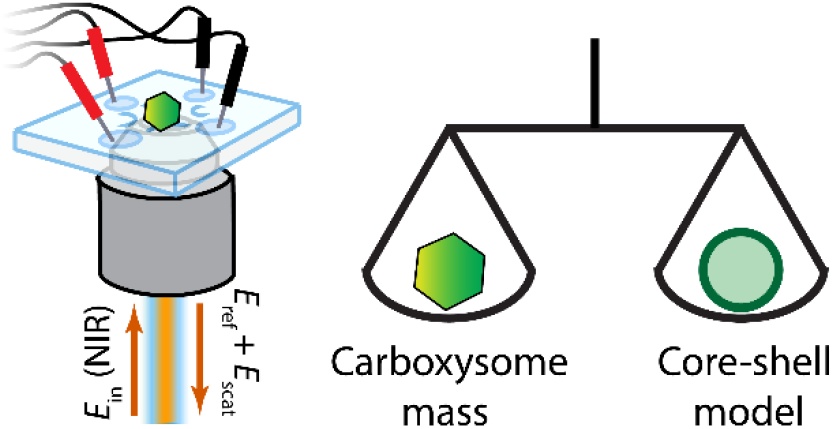

## Introduction

Biological processes use compartments at the nanoscale to enclose particular materials and maintain distinct environments. While some of these compartments exist primarily to transport chemicals and biomolecules outside the cellular environment (e.g. viruses, extracellular vesicles, etc.), others maintain distinct environments for chemical reactions (e.g. lysosomes, peroxisomes, etc.).^1^, ^2^ In prokaryotes, a variety of microcompartments have been described, and we focus here on the carboxysome – a key component of the carbon-concentrating mechanism of many autotrophic bacteria.^3^, ^4^ This intracellular compartment encloses the enzyme rubisco alongside other proteins^5^ and creates a chemical environment to increase the efficiency of fixing carbon dioxide to organic carbon,^6^ estimated to contribute 10-25% of annual global carbon fixation.^7^ Since carboxysomes spontaneously self-assemble, they can significantly vary in size, shell integrity, and enzyme concentrations and can be programmed for engineering purposes.^8^ Therefore, there is a need to understand the composition and chemistry of carboxysomes not only in their average properties, but also in terms of their heterogeneity and dynamics. In particular, to analyze the composition of carboxysomes it would be valuable to measure not only the total external size of the shell but also the contents of the interior at a single-carboxysome level.

The mass of microcompartments is a good place to start. A number of approaches have been developed to determine the mass of single tiny objects. Mechanical oscillator^9^ and cantilever-based^10^, ^11^ methods have been developed for directly weighing individual nanoparticles. Measuring mass and structural information about proteinaceous assemblies has also been achieved with scanning transmission electron microscopy (STEM)^12^ and cryo-Electron Tomography (cryo-ET) more recently.^13^ While these techniques all report on mass, they are potentially destructive or can be difficult to combine with other reporting modalities, limiting the information that can be obtained for each particle.

Among nondestructive approaches, total mass can also be assessed by first measuring light scattering, which directly senses refractive index. Then mass can be inferred if the material class (protein or DNA) of a particle is known. This can range from optical tweezers-based force constant measurements to side-scatter in flow cytometry.^14^, ^15^ In particular, a relatively new approach uses interferometric scattering microscopy^16^, ^17^ (iSCAT) to measure the mass of single proteins which land on a surface down to 10s of kDa, called “mass photometry”.^18^ This technique offers high mass resolution for objects at a well-defined optical interface,^19^, ^20^ but solution-phase objects are more challenging to measure due to their rapid diffusion and the change in the particle’s point-spread function with axial position.^21^ A recent study combines the single particle tracking of iSCAT with the refractive index sensitivity to measure the densities of vesicles using single-particle trajectories on the order of 100s of ms.^22^ For longer timescale biological dynamics (> 1s), there remains a need for sensitive, nondestructive single-particle mass measurements of protein assemblies in the liquid phase compatible with fluorescence reporting schemes, for example. Fluorescence can enable sensing of additional chemical aspects of the particle,^23^ and can even allow measurements of the mass distribution in the particles, as we show in this work.

One approach to increasing the observation time of single particles in free solution for spectroscopy is to apply active feedback to cancel out the Brownian motion – what we call anti-Brownian Electrokinetic (ABEL) trapping.^24^, ^25^ Using native or added fluorescence from the single object, ABEL traps have held single molecules and protein assemblies in solution for many seconds to directly monitor their chemical dynamics, assembly, and photophysics.^26–29^ More recently, we have used interferometric scattering to detect the position of a particle for applying feedback, creating the Interferometric Scattering Anti-Brownian Electrokinetic (ISABEL) trap.^30^ In recent advances to the technique we have shifted the high intensity scattering illumination laser to the near-infrared (NIR) and implemented simultaneous interleaved fluorescence excitation and detection of reporters in the visible to yield a more flexible ISABEL trap.^31^ This instrument was able to trap single carboxysomes and precisely measure their redox states from fluorescence excitation spectroscopy over ~ 1s.

While the new ISABEL trap enables ratiometric fluorescence reporting, in this work we use the scattering signal to determine the mass of each trapped object in solution. With a calibration using standardized bead samples, we obtain scattering cross-sections for each trapped object. Using this scattering cross-section along with a core-shell Mie scattering model and the knowledge of refractive indices expected for proteins, we extract the masses of individual carboxysomes in solution. Furthermore, we use simultaneous signals from fluorescent protein constructs targeted to the carboxysome cargo to probe the interior distribution, estimating the amount of core material inside each carboxysome. By analyzing the scaling relationship between the cargo loading and the scattering from the whole object, we are able to estimate both the total mass and the cargo density distribution inside carboxysomes.

## Materials and Methods

### Polystyrene nanoparticles

NIST-traceable polystyrene nanospheres of nominal diameter 80 nm were obtained from Thermo Scientific (Cat. No. 3080A, lot no 242108) with certified mean diameter 81 nm ± 3 nm, (95% confidence interval) at approximately 1% solids, and diluted 10,000x in nanopure water (Easypure UV/UF) for trapping experiments. Fluosphere carboxylate-modified microspheres with yellow-green fluorescence (505/515 absorption/emission peak wavelengths) and 0.1 μm nominal diameter were obtained from Invitrogen (Cat. No. F8803, Lot no. 1588588) at 1% solids and diluted 40,000x in nanopure water for trapping experiments. Carboxyl-modified polystyrene microspheres of nominal diameter 0.135 μm were obtained from Bangs Laboratories, Inc (Cat. No. PC02005, Lot no. 15033) at 10% solids and diluted 80,000x in nanopure water for trapping experiments.

Polystyrene nanoparticle diameters were determined from cryoTEM images by manually selecting points on their boundary and fitting circles to the data. The nominally 0.1 μm fluorescent polystyrene nanospheres appear normally distributed with mean μ = 108 and standard deviation σ = 7 nm (Fig S1). The nominally 0.135 μm polystyrene nanospheres appear normally distributed with μ = 130 and σ = 2.4 nm (Fig S2), summarized in Table S1.

### Ultra-uniform gold nanoparticles

Two samples of ultra uniform gold nanospheres capped with PEG_12_-carboxylate functional groups were obtained from Nanocomposix, with nominal diameters 30 nm (Cat. No. AUXU30, Lot. No. JSF0124) and 50 nm (Cat. No. AUXU50, Lot. No. SDC0158) at 0.05 mg/mL of gold. The actual mean and standard deviations of diameters reported by the manufacturer, calibrated from TEM, were 28.0 ± 0.9 nm and 51.0 ± 0.9 nm (see Table S1). The nanospheres were diluted 50x and 6x in nanopure water for trapping experiments.

### Synthesizing and purifying carboxysomes

Carboxysomes containing Superfolder GFP (sfGFP) were expressed heterologously in *E. coli BW25113* bearing two plasmids: one with the *H. neapolitanus* HnCB10 carboxysome operon,^32^ and the other with sfGFP fused to the first 53 residues of the carboxysomal carbonic anhydrase, CsoSCA. This CsoSCA fragment has been shown to interact with rubisco and enables the targeting of foreign cargo into the carboxysome interior.^33^ The HnCB10 operon was modified with an N-terminal FLAG tag on the hexameric shell protein CsoS1A.

Carboxysomes were expressed by growing the cells to mid-log phase in 100 mL LB media at 37°C. At OD_600_ of 0.6-0.8 the temperature was reduced to 18°C and protein expression induced with 1mM IPTG and 100 nM aTc. Cells were grown overnight before pelleting and freezing at −20°C for storage. Carboxysomes were purified by lysing the cell pellets with 2 mL B-PER (Thermo Scientific) supplemented with 10mM MgCl_2_, 20mM NaHCO_3_, 1mM PMSF, 0.01 mg/mL DNaseI, and 0.1 mg/mL lysozyme. After 1 hour incubation at room temperature the lysate was clarified with a 12,000 × g spin for 20 min. The clarified lysate was centrifuged for 30 min. at 40,000 × g to pellet the carboxysomes. The carboxysomes were gently resuspended in 2 mL TEMB buffer (10 mM Tris-HCl, 10 mM MgCl_2_, 20 mM NaHCO_3_, 1 mM EDTA, pH 8) and centrifuged again at 40,000 × g. Pelleted carboxysomes were gently resuspended with 0.2 mL of TEMB on ice and then loaded onto a 3-mL 10-50% sucrose (w/v) gradient in TEMB. The gradient was centrifuged for 18 min. at 105,000 × g then fractionated into 0.2 mL fractions and analyzed by SDS-PAGE. The carboxysome containing fractions were pooled and centrifuged for 1 hour at 105,000 × g to pellet carboxysomes. Carboxysomes were resuspended in TEMB and stored at 4°C. Past experience has shown that carboxysomes stored at 4°C retain morphological integrity for months as visualized by negative stain TEM. All samples analyzed in this study were measured within six months of purification.

### Carboxysome radius determination

Effective diameters were determined from cryo-TEM images (details below) by measuring the area of the polygon manually traced around the carboxysome shells, then evaluated as d_eff_ = (4A/π)^1/2^ assuming a circular shape. As shown in Fig. 5e, the diameters appear normally distributed with μ = 112 nm and σ = 17 nm.

### Preparing carboxysomes for trapping

For trapping experiments, carboxysomes were drawn from the top of the stock suspension so as not to retrieve any large aggregates that had precipitated out of suspension. Carboxysomes were diluted into HEPES buffer (10 mM HEPES, 15 mM NaCl, pH 7.5, 0.2 μm filtered) such that less than one carboxysome would be in the trapping volume at any given time, roughly 1-10 pM. The carboxysomes were labeled in most situations with anti-FLAG antibodies conjugated to Alexa Fluor 647 (Invitrogen MA1-142-A647), but this antibody fluorescence is not used in the present analysis. Briefly, 1 μL of the carboxysome stock was incubated with 1 μL of antibodies diluted to 0.14 mg/mL in HEPES buffer for 5 hours. The mixture was diluted to 200 μL with HEPES buffer immediately before trapping.

### Cryo-electron microscopy of samples

3 μL of carboxysome suspension (diluted 10x in HEPES buffer from stock) or polystyrene nanosphere solution was deposited onto a glow-discharged holey carbon electron microscopy grid (Quantafoil R 2/2 G200F1), blotted on both sides for 2 to 3 seconds, and plunge frozen into liquid ethane (Gatan CP3). Cryo-electron transmission micrographs were acquired on a 200-keV electron microscope (Thermo Fisher Glacios) equipped with a direct detector (Gatan K2). Images were acquired with pixel spacing of 2.43 Å with defocus targets for carboxysomes and polystyrene nanospheres of −5 μm and −10 to −1.5 μm, respectively.

### Calculating scattering cross-sections with Mie theory

The scattering cross-sections for uniform spherical particles and spherical core-shell particles were calculated using the Mie scattering approach described in Bohren and Huffman,^34^ implemented in Matlab.^35^ All refractive indices were utilized at 802 nm and 20 C. The refractive indices used for polystyrene and gold were 1.5781^36^, ^37^ and 0.15413 + 4.9243i^37^, ^38^ respectively. The protein densities were converted to refractive index using a literature refractive index increment of dn/dc = 0.1847^39^, converted to 802 nm^40^.

The scattering cross-sections for all the polystyrene beads were calculated assuming uniform spheres. The mean and uncertainty for the scattering cross-section of the 80 nm polystyrene nanospheres were calculated as the mean and standard deviation from 1000 samples of diameters from a Gaussian distribution of 81.0 ± 1.5 nm, the manufacturer quoted 1σ uncertainty on mean diameter (Fig S3b). The mean and uncertainty for the scattering cross-sections for the two sets of gold nanospheres were calculated in two steps using manufacturer provided diameters of 28.0 ± 0.9 nm and 51.0 ± 1.9 nm. As each of the gold beads were PEG-carboxylate capped, we adjusted the scattering cross-section for surface coating as follows. We sampled Au-PEG core-shell models with a random number of PEG_12_-COOH/nm^2^ blocking groups on the gold surface between 3 and 5.^41^ The coating density is much less precisely known than the diameter, and so the scattering cross-sections were averaged for 1000 sampled Au sphere diameters for a specific coating density, and then the mean and standard deviation of 100 samples with different coating densities were calculated (Fig S3a and S3c respectively). We assumed a PEG shell of thickness 4.68 nm, each molecule of weight 617 Da, and a refractive index increment proportional to density in this shell. The refractive index used for PEG was 1.461 for a density of 1.1257 g/mL^42^, corrected for 802 nm.

### Microfluidic trapping cell

The trapping experiments were performed in a quartz microfluidic cell that has been previously used for ABEL and ISABEL trap experiments and has been described in more detail previously.^30^, ^43^, ^44^ The cells consist of two crossed channels ~10 μm deep etched into a quartz piece that feature a shallower ~1.5-2 μm thin trapping region at the center (Figure 1a), with four ports to load solutions and to admit platinum electrodes for feedback forces in × and y. The bottom of the cell is formed by a chemically bonded 0.15 mm thick quartz coverslip. The cells are securely held in a custom-machined holder to minimize mechanical drift contributing to noise in the interference pattern.

**Figure 1.**
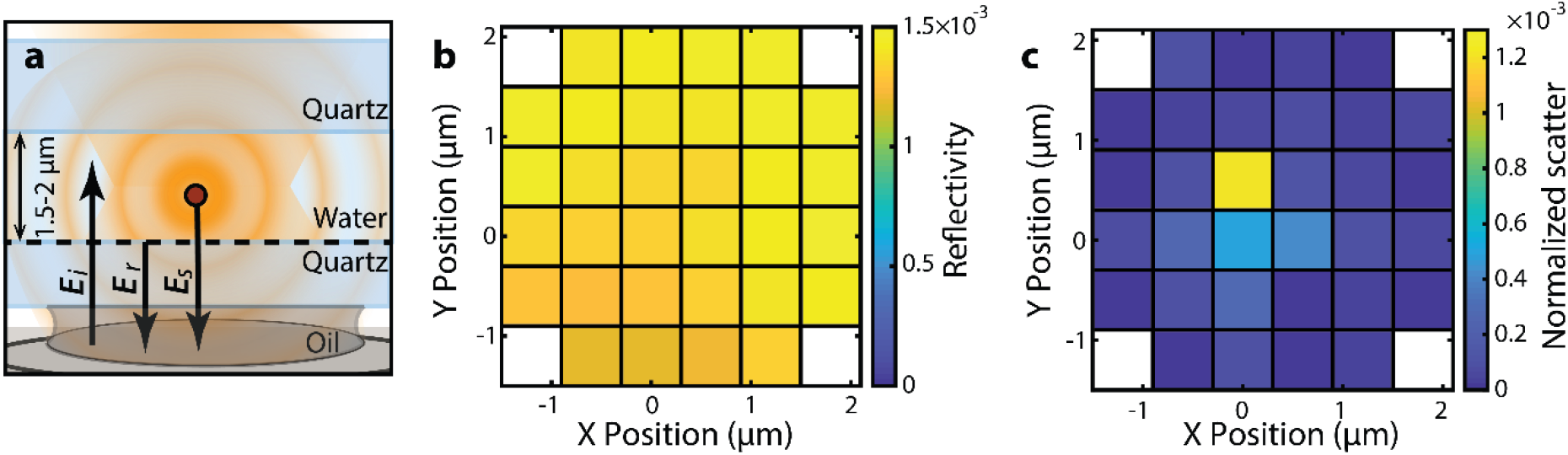
Measuring the scattered field in the microfluidic trapping cell. (a) A side-view schematic of the central trapping region of the microfluidic cell. The incoming focused NIR field *E*_i_ gives rise to the reflected field *E*_r_ from the quartz-water interface, and the scattered field *E*_s_ from a scattering object. (b) The reflectivity of the interface varies over the 32 beam-scan positions between 1.2 × 10^−3^ and 1.6 × 10^−3^. (c) The scattering contrast normalized by reflectivity shows increased signal at the position of the scattering object for a 0.1 μm fluorescent polystyrene nanosphere.

The cells are chemically treated before each experiment to prevent sticking. For the polystyrene and gold nanoparticle trapping experiments, the cells are treated with 1M KOH for ~ 15 minutes to create a negative charge at the quartz interface^30^. For the carboxysome trapping experiments, the cells are passivated with a polyelectrolyte multilayer consisting of four alternating layers of poly(ethyleneimine) and poly(acrylic acid) resulting in an anionic surface^31^.

### ISABEL trap optics

The ISABEL trap experiments were performed on a custom-built optical setup as has been recently described^31^ (see also Note S1 and Fig S4). Briefly, a near-IR (NIR) laser at 802 nm is scanned to different x-y positions in the sample plane with a pair of acousto-optic deflectors (AODs) in a 32-point Knight’s tour pattern. The NIR spot at the sample was ~ 500 nm in diameter, and the pitch of the scan pattern was chosen to provide a time-averaged uniform intensity of ~ 250 kW/cm^2^ over a 3.6 μm × 3.6 μm region.^31^ The back-scattered and reflected light were separated from the pumping beam using a polarizing beam-splitter and quarter-wave plate combination and focused through an iris to block unwanted light onto a large area photodiode (Newport 2031) for interferometric scattering detection. The fluorescence excitation laser at 488 nm follows a path distinct from the AODs and is aligned to overlap with the NIR beam scan pattern using a 775SP dichroic. This beam illuminates a Gaussian spot at the sample with 1/e^2^ radius of 5.5 μm, which covers the NIR Knight’s tour pattern. The fluorescence collected in the backward direction is separated by a 405/488/561/647 quad-pass dichroic, and is passed through a pinhole equivalent to 3.6 μm diameter in sample space. Green fluorescence (500-560 nm) is collected on a Si single-photon avalanche photodiode (τ-SPAD, PicoQuant).

### ISABEL trap control and feedback scheme

The ISBAEL trap hardware is controlled primarily by a field-programmable gate array (FPGA) on a National Instruments card (RIO PCIe-7856R). The NIR beam is positioned at each spot in the scan pattern for 18.75 μs, and the photodiode signals from the corresponding dwell times are used to construct a single “frame” composed of all 32 points over 600 μs. The background is estimated by averaging frames over 10ms when the feedback is off and no object is transiting through the trap. Each frame for trapping is normalized by the NIR beam power, and the absolute value of the fractional change from background |(*P*_*det*_ − *P*_*bkg*_)/*P*_*bkg*_| is calculated for each position of the NIR beam. Feedback force is applied proportional to the displacement from trap center of the position of greatest absolute fractional change at the end of every 600-μs period by the amplified FPGA.^31^ The FPGA also records arrival times of photons on the SPAD channel with 12.5 ns resolution.

## Results

### Normalizing scatter for differences in reflected power

The fractional interferometric signal above background described above is sufficient to obtain a useful real-time signal for trapping, but this quantity changes with the varying reflectivities of different trapping cells and beam position in the trapping cell. We now describe how the signals from the ISABEL trap can be used to extract quantitative scattering information about the object in the trap. The interferometric signals on the NIR detection photodiode is spatially filtered to the ~ 5×5 μm^2^ around the primary reflection with an iris approximately conjugate to the focal plane to reject unwanted reflections from other interfaces in the optical path. The sample z-position is reproducibly set to obtain a sharp image of the reflection of the illumination scan pattern. The vertical alignment of the scattering illumination beam is checked before each day of experiments, and the magnitude of the reflected power from a test glass coverslip-air interface is measured for each point in the scan pattern, 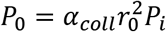, where *α*_*coll*_ is the efficiency of transmission of the reflected beam to the detector, 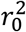 is the computed Fresnel reflectivity of the glass-air interface (0.040) for near-normal incidence, and *P*_*i*_ is the incident power. This glass-air reflectivity measurement is essential to normalize the intensity of the local oscillator beam during detection.

When a scattering object enters the trap region, the electric field at the detector results from the interference of the scattered field *E*_*s*_ from the object and the total reflected field *E*_*r*_ = *r E*_*i*_ from the quartz-water interfaces (Fig. 1a). While there are two potential sources of reflected fields, one is out of focus and is largely blocked by the iris before detection. The field reflectivity coefficient *r* depends on surface variations and residual bonding material at the quartz coverslip-water interface, and it can thus vary from position to position in the trapping region. The scattered field varies in space and is a function of the incident field at the position of the particle as well as the strength of the scattering response from the particle, but it will be necessary to carefully analyze the situation to obtain quantitative results for the scattering strength.

We are now in a position to clarify the various signals that can be measured and how they relate to the strength of the scattering. In dark-field scattering, the strength of the scattering response can be characterized in terms of the scattered power that reaches the detector *P*_*scat*_ = *α*_*coll*_*s*^2^*P*_*i*_ with *s* the dimensionless field scattering response. Further, the quantity we seek, the scattering cross-section *σ_scat_*, is defined by *P*_*scat*_ = *α*_*coll*_*σ*_*scat*_*I*_*inc*_ with *I_inc_* the intensity at the scatterer, hence *s*^2^ ∝ *σ*_*scat*_. With both the reflected and scattered fields important in the ISABEL trap, the total power at the detector includes interference between the two fields and can be expressed as

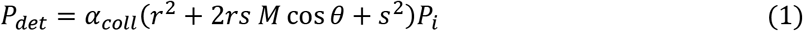

where *M* is a factor accounting for the mode overlap between the scattered and reflected fields, and *θ* is the phase difference between the two. The pure scattering term *s*^2^ is generally negligible compared to the interferometric scattering cross-term. The pixel-by-pixel background for a trapping experiment is estimated during post-processing as the 100-frame moving median of the detected power where the fluctuations of phase force the cross term to have a median contribution of zero, *P*_*bkg*_ = *α*_*coll*_*r*^2^*P*_*inc*_. The running median accounts for slow variations in the reflected field such as fringes moving across the pattern. The background is further corrected for power fluctuations on a frame-by-frame level using the average reading from the outer 16 pixels as described previously^30^. The scattering contrast 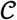 is calculated at each scan position as the fractional difference from the background power,

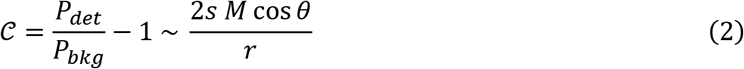

analogous to iSCAT analysis^18^. However, the reflection coefficient *r* varies across space in an ABEL trap cell and must be calculated for each scan position by comparison to the reflected power *P*_0_ from the test glass coverslip-air interface, with 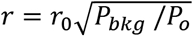. An example of a map of the reflectivity for a trapping experiment is shown in Figure 1b, with a ~30% difference between the largest and smallest reflectivities. Correcting for differences in reflectivity at different positions then gives us the normalized scatter

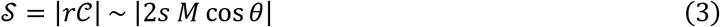

which should depend only on the scattering cross-section of the object and the relative position and phase. The normalized scatter values for a single frame of trapping data (single Knight’s tour) for a 0.1 μm fluorescent polystyrene nanosphere are shown in Figure 1c. The position of the nanosphere is visible as the position of greatest normalized scatter. The reader will recognize that the trapped object can move in *z*, which changes the phase factor in Eqn (3). To address this, objects are trapped over times longer than the z-motion fluctuations, and the time average of the normalized scatter is used to yield quantities of interest without confusion from the phase factor (Fig S5). This time averaging gives a normalized scatter measurement 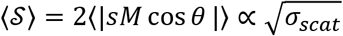, which, with proper calibration, enables extraction of the single-particle scattering cross-section as shown in Fig. 3 below.

**Figure 2.**
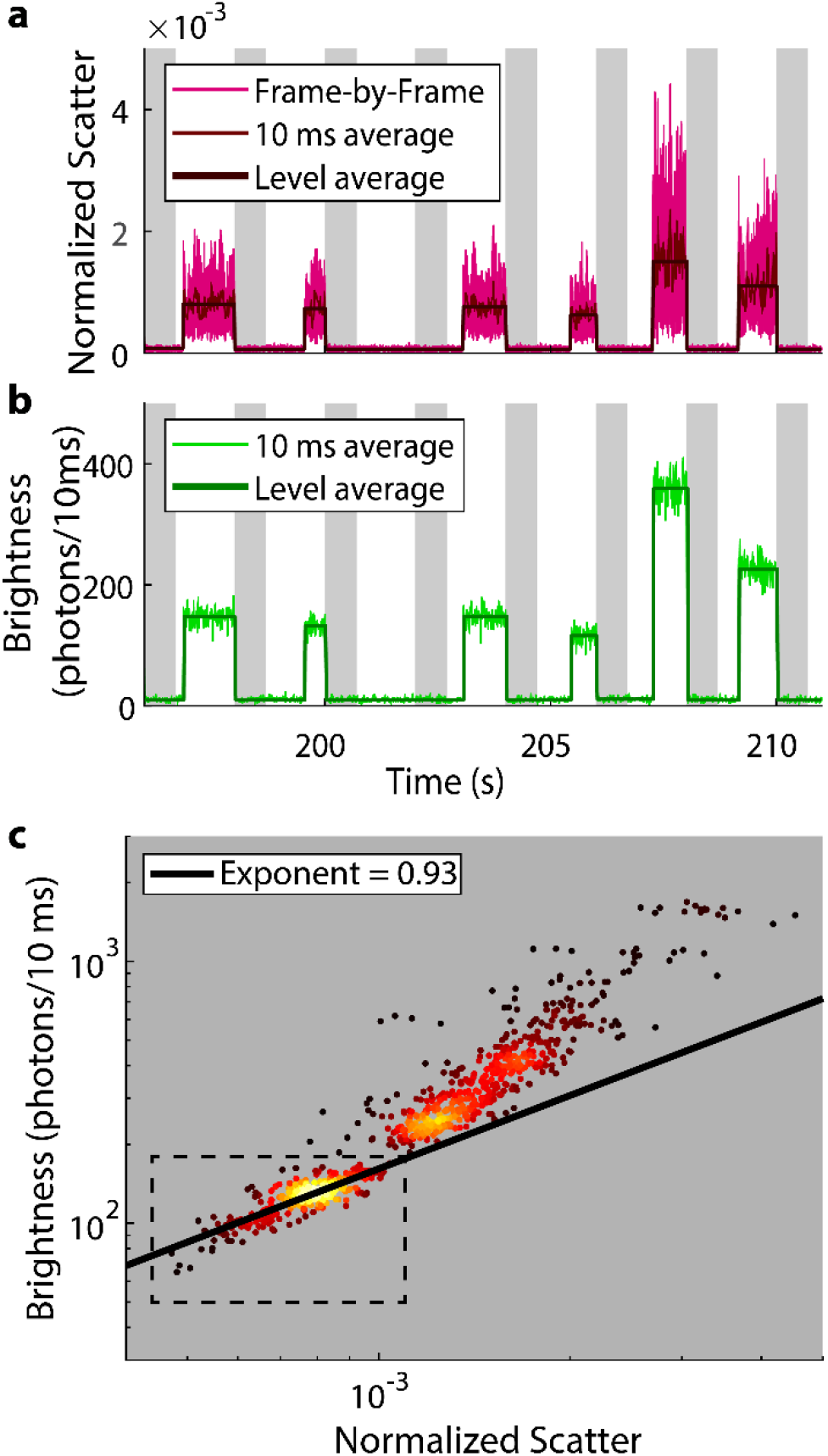
Trapping 0.1 μm fluorescent polystyrene nanospheres. (a) A time trace of the normalized scatter, for each 32-point scan or “frame” (light red, trace with greatest fluctuations), averaged over 10 ms (darker red), and averaged over an entire level from level finding (Black line). Grey background represents the time during which the feedback is turned off. (b) The fluorescence brightness for the same time trace, averaged over 10 ms (light green) and an entire event (dark green). (c) The fluorescence brightness *vs* normalized scatter distribution of all the levels observed as trapping events in log space. Each point is colored by the local density of neighbors. A power law fit to the lower population in the box (putative monomers, N = 318) yields an exponent of 0.93, suggesting volume loading of fluorophores in the nanospheres.

**Figure 3.**
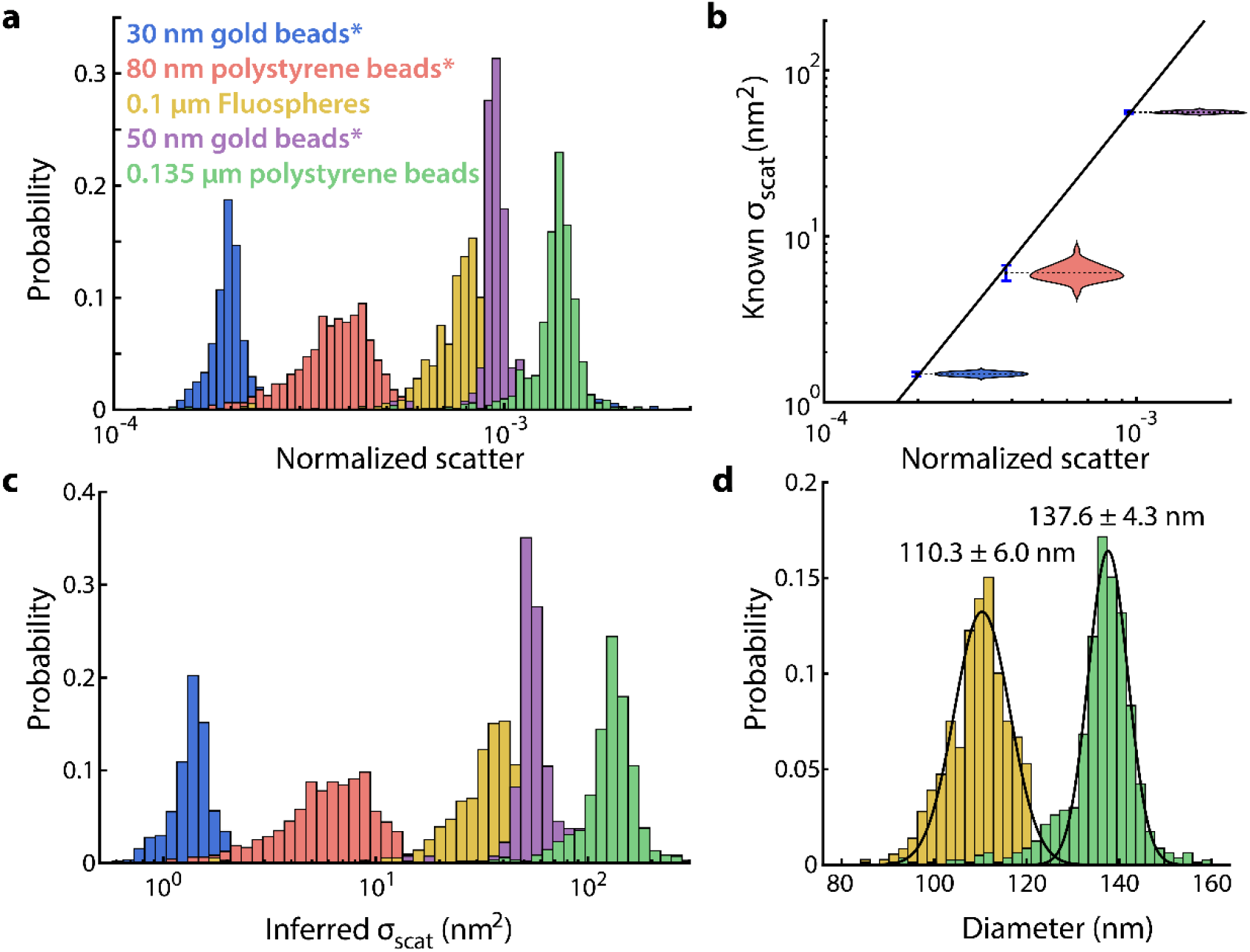
Converting measured scatter to scattering cross-sections for nanospheres. (a) Measured normalized scatter levels for 3 sizes of polystyrene nanospheres and two sizes of gold nanospheres, histograms on log axes (N = 1083, 949, 359, 134, 819 from left to right). The numbers in the legend correspond to nominal diameters, and the nanosphere samples used to calculate the conversion from normalized scatter to scattering cross-section are marked by *. (b) The extracted relation (Equation 4) between scattering cross-section and normalized scatter from fitting a power law to the mean calculated Mie scattering cross-section vs mean normalized scatter for the three calibration sets of nanopheres. The error bars are S.E.M. corresponding to the distribution of calculated Mie scattering cross-section (violin plots, displaced laterally) from manufacturer quoted diameters. (c) The scattering cross-sections inferred from the measured normalized scatter for each of the samples in (a), on log axes. (d) The diameters for the 0.1 μm fluorescent polystyrene nanospheres and the 135 nm polystyrene nanospheres, deduced from the inferred scattering cross-sections based on a Mie scattering model. The lines denote Gaussian fits to the histograms.

### Trapping fluorescent polystyrene nanoparticles

Since the goal of this paper is to connect scattering measurements in the ISABEL trap to simultaneous fluorescent assessments of the trapped object, we now describe how the two measurements can be combined. Fluorescent polystyrene nanoparticles of 0.1 μm nominal diameter provide a useful control sample as spheres of constant refractive index and uniform fluorescence. Data from trapping fluorescent polystyrene nanospheres with fluorescence emission peaked at 515 nm are shown in Figure 2. The maximum normalized scatter over the center 3 × 3 positions (SI note) in the trapping pattern for a representative portion of the trapping trace is shown in Figure 2a. When feedback is enabled, fluorescent nanoparticles arrive in the illuminated region of the microfluidic cell and are trapped, giving a normalized scatter value that fluctuates between two extrema (~2 × 10^−4^ to ~2 × 10^−3^ for the first trapping event in Figure 2a) consistent with a single average scattering strength from frame to frame, where the fluctuations arise from changes in the interferometric mode overlap and the relative phases of the scattered and reflected fields. The 10 ms binned signal (darker line) shows a more constant level. Trapping events can be identified with a level-finding algorithm on this average (Note S2), and the normalized scatter averaged over a level is plotted as well (Fig 2a, darkest line). The feedback is toggled on and off regularly to release trapped nanoparticles, catch new nanoparticles, and collect statistics on many single nanoparticles. The levels return to background when feedback is off, ensuring that the objects are not stuck to the microfluidic cell surfaces. The fluorescence photon arrivals recorded at the same time, binned every 10 ms, are plotted in Figure 2b. The fluorescence shows increases from background at the same time as increased normalized scatter, confirming that the trapped objects are the fluorescent polystyrene nanospheres. The fluorescence can also be averaged over the time periods identified by level-finding on the normalized scatter (darker green in Fig 2b). This allows each individual particle to be characterized in terms of its average normalized scatter and average fluorescence brightness.

The distribution of normalized scatter and fluorescence levels from fluorescent polystyrene nanospheres is plotted as a scatter plot with coloring based on numbers of nearest neighbors^45^ in Figure 2c. Because the data span orders of magnitude, we plot the levels on logarithmic axes for ease of visualization. At least two distinct populations can be discerned, one peaked approximately at a normalized scatter of 8 × 10^−4^ and a fluorescence brightness of 140 photons/10ms and the other broadly extending up from normalized scatter 1×10^−3^ and brightness 200 photons/10ms. The lower population is expected to arise from monomers, while the broad higher population from dimers and perhaps larger aggregates. The fluorescence increases with the normalized scatter, likely due to increases in the nanosphere diameter leading to a larger number of loaded fluorophores and an increased scattering cross-section, respectively, with additional variations in this trend arising from heterogeneity of fluorescent label loading in the nanospheres and noise in the measurement of both variables (Note S2). This increase of fluorescence with scatter can be characterized by fitting a power law (black line in Fig 2c) to the population of monomers (in the dashed box), which yields an exponent of 0.93 ± 0.06, consistent with volume loading of fluorophores being proportional to the mass. The deviation of the power law from near unity for the larger particles is likely due to nonspherical shapes of the aggregates. The same principles involved in identifying particles of interest in a population and testing the trend of fluorescence brightness vs scatter against a model can be taken from relatively uniform particles and applied to particles with more complicated composition such as carboxysomes, *vide infra*.

### Converting measured scatter levels to scattering cross-sections

The relation between the average normalized scatter for a trapping event and the scattering cross-section of the trapped object can be calibrated by trapping different nanoparticles of known sizes. We will use measurements of three highly-calibrated bead samples to make this connection, and then show that the resulting relationship can be used to predict the behavior of two other bead samples. The histograms of normalized scatter values averaged over each trapping event for five different samples are shown in Figure 3a. For each sample, only the values close to the first peak of the normalized scatter were chosen for further analysis and are plotted, to avoid dimers and outliers analogously to Fig 2c. The nominal 30 nm and 50 nm gold nanospheres, and the 80 nm polystyrene nanospheres are well-characterized in their diameters for use as electron microscopy calibration standards, and thus have the most accurately and precisely known mean values of their scattering cross-section (Note S3). Consequently, these three samples are shown with an asterisk in the figure. The spreads in the calculated scattering cross-sections are shown in Figure 3b (violin plots).

The normalized scatter at any given time for an object in the ISABEL trap depends not only on the scattering cross-section of the object, but also on its position relative to the incident beam. However, over an entire trapping level we expect the object to explore the full space of positions such that the average trapping level depends only on the scattering cross-section. The inverse relation can then be expected to be 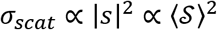. By fitting a power law to the mean scattering cross-section vs mean normalized scatter of these samples as shown in Figure 3b (Note S3), we can determine a functional form for the scattering cross-section of the form

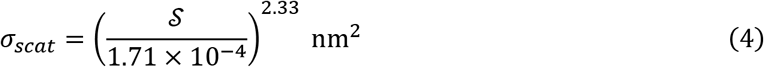

in terms of the average normalized scatter signal. We see a deviation from the expected exponent of 2 by ~15%, in a direction consistent with increased measured scattering cross-sections from the maximum-of-absolute-value calculation (Note S3, Fig S6). The normalized scatter for all the samples measured in Figure 3a can be converted to single-particle scattering cross-sections using Eq. (4), as plotted in Figure 3c. The measured scattering cross-sections span two orders of magnitude from 1 nm^2^ to over 100 nm^2^, with a spread for each sample.

To validate our calibration, we use Eqn (4) and the normalized scatter measurements from the 100 nm and 135 nm nanosphere samples not used in the calibration to infer their scattering cross-sections and, therefore, their diameters. For these polystyrene nanospheres with a uniform composition and known refractive index, the scattering cross-section for any given diameter value can be calculated from Mie scattering theory. Using this, the measured scattering cross-sections can be converted to single-particle diameters as shown in Figure 3d. The nominally 100 nm and 135 nm polystyrene nanospheres when fit with a Gaussian distribution show a mean diameter and standard deviation of 110.3 ± 6.0 nm and 137.6 ± 4.3 nm respectively, consistent with the manufacturer quoted values of 0.1 ± 0.0063 μm and 0.135 μm, which is also consistent with diameters we measured by cryo-TEM of 108 ± 6 nm and 130 ± 2.4 nm. The calibration in Eqn (4) above was extracted from objects in the normalized scatter range most relevant to our measurements on carboxysomes (see Figure 4e below), and has been tested on objects in the relevant size range. The good match between cryo-TEM measured diameters and those inferred from ISABEL trap measurements on nanoparticles not used in the calibration (Eqn 4) increases the confidence in our single-particle scattering cross-section measurements as applied below to biological samples.

**Figure 4.**
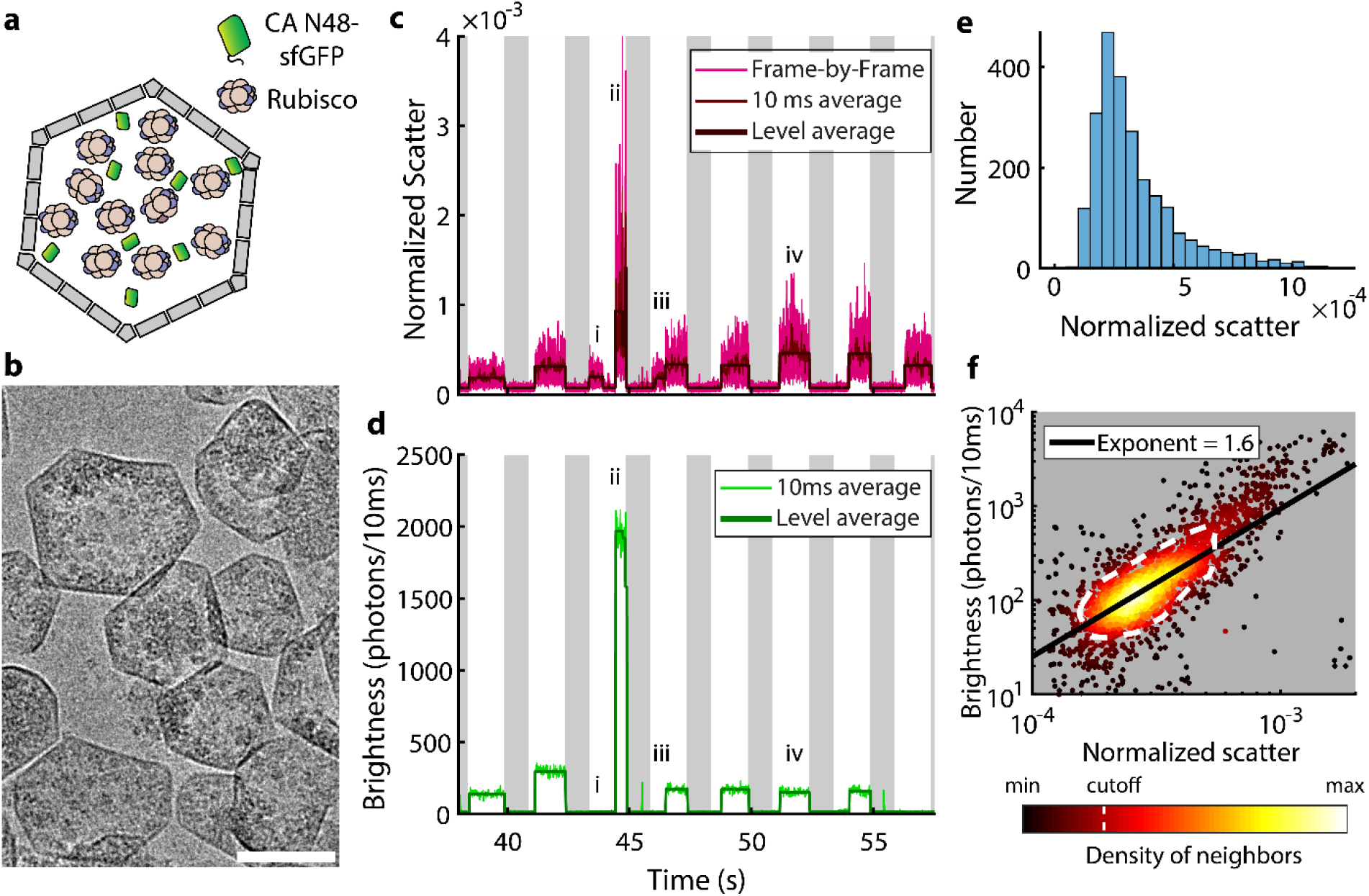
Trapping carboxysomes with scattering combined with simultaneous fluorescence measurements. (a) A schematic of the carboxysomes expressed in *E. coli*. Super-folder GFP is targeted to the carboxysome interior with a sequence expected to bind rubisco. (b) Cryo-TEM images of the purified carboxysomes. Scale bar: 100 nm. (c) Time trace of the normalized scatter from carboxysomes in the ISABEL trap, with levels identified as above. (d) Time trace of the sfGFP fluorescence brightness for the same trapping data as (c). Events marked i-iv in both channels are described in the text. (e) Histogram of normalized scatter values for carboxysome trapping levels with GFP fluorescence above background. (f) A scatterplot of fluorescence brightness vs normalized scatter on log-log axes with points colored by density of neighbors. The power law line to characterize the shape is fit to the points above a density threshold, roughly indicated by the white dashed line.

**Figure 5.**
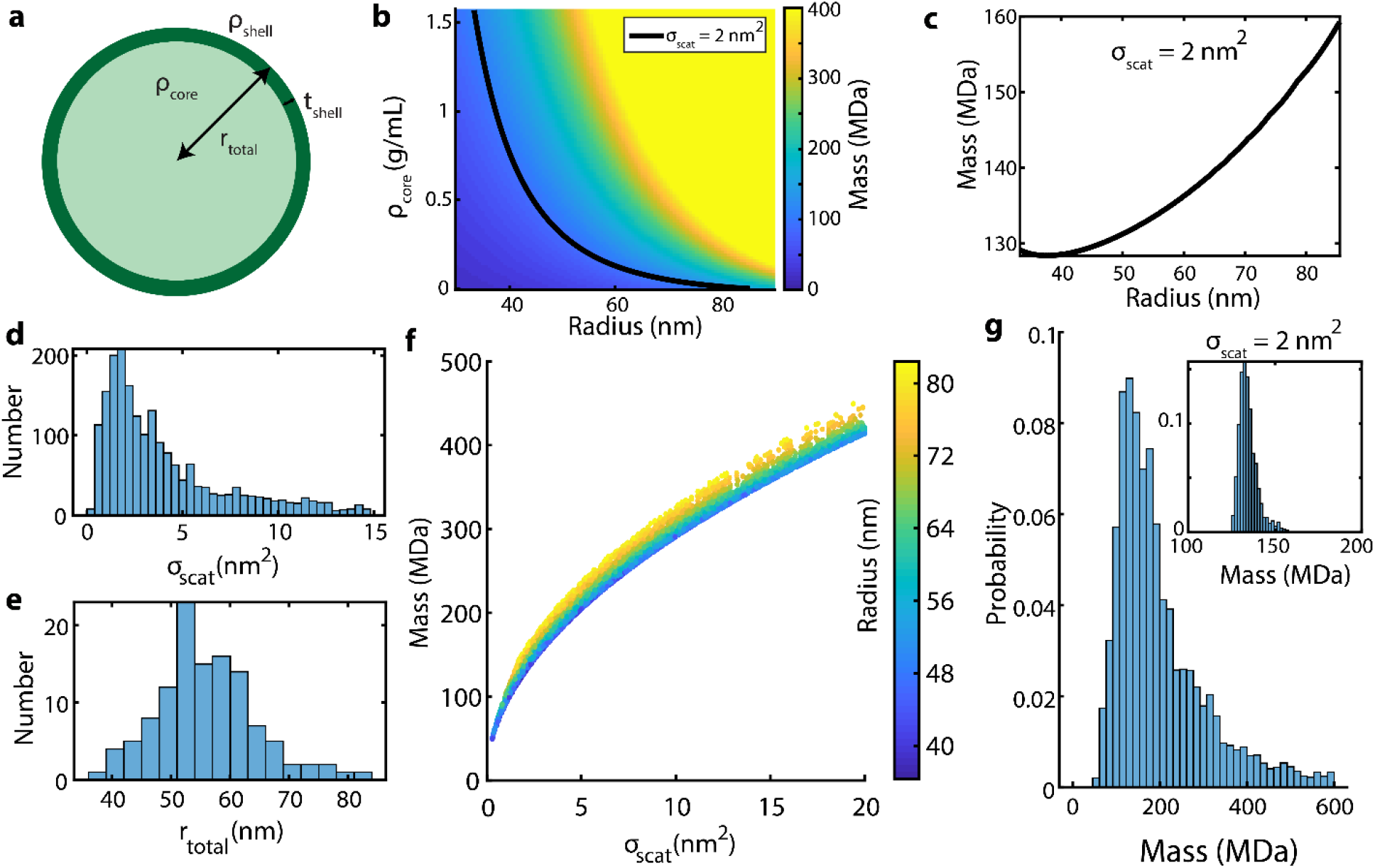
Calculating carboxysome masses. (a) A spherical core-shell model of a carboxysome for Mie scattering analysis. The shell thickness and density are assumed to be known. (b) The calculated mass of the carboxysome (colormap) for different values of core density and total radius. The line represents carboxysomes with *σ*_*scat*_ = 2 *nm*^2^. (c) The mass along the *σ*_*scat*_ = 2 *nm*^2^ line. (d) The distribution of scattering cross-sections measured for the carboxysomes from Fig. 4e. (e) Distribution of radii measured by cryo-TEM. (f) The distribution of calculated masses vs measured scattering cross-sections, colored by the radius. (g) The histogram of calculated masses, with the mass distribution for a single scattering cross-section inset.

### Moving to biological samples: Trapping single carboxysomes labeled with sfGFP

Using our studies of single component nanoparticles as a foundation, we are now in a position to address the far more complicated situation when a real biological nano-object is studied in the ISABEL trap. Carboxysomes offer an interesting example of a nanoparticle that may not have a uniform composition or filling of its interior. We seek to use the scattering and the additional fluorescence variable to infer more information about these objects. α-Carboxysomes from *H. neapolitanus* were expressed heterologously in *E. coli*, together with sfGFP fused with a targeting sequence (see Methods) to direct the sfGFP to the interior of the carboxysome (Figure 4a) via noncovalent binding to rubisco. A cryo-TEM image of purified carboxysomes is shown in Figure 4b. The carboxysome shells (darker lines) are roughly hexagonal in the 2D projection, suggestive of the icosahedral shapes expected from prior studies.^13^, ^46^ However, the shapes are nevertheless irregular suggesting deviations from the canonical carboxysome structures, previously seen for α-carboxysomes grown in *E. coli*.^32^ The sizes of the carboxysomes vary substantially, with most in the ~100 nm diameter range. The rubisco units inside the carboxysomes are also visible as dark near-circular shapes on the order ~10 nm. The carboxysomes vary greatly in the number of rubisco complexes inside, suggesting a large variability in size and filling which we show can be investigated in the ISABEL trap.

A time trace of carboxysome trapping is shown in Figure 4 c, d, showing normalized scatter signal and corresponding fluorescence brightness, respectively. The carboxysomes show clear levels that end when feedback is disabled, indicating successful trapping events with negligible sticking. As with the beads, we deliberately alternated feedback on and off in order to efficiently sample the wide distribution of carboxysomes, but we note that longer trapping times up to minutes is feasible for single particles.^30^, ^31^ The normalized scatter displays fluctuations expected around each level from position changes, similar to what was observed for polystyrene beads in Figure 2a. The scatter and the fluorescence levels both span large ranges. Certain events such as i and iii show increased scatter levels above background, but no corresponding increases in the fluorescence. This suggests that the objects trapped at these times do not have sfGFP molecules, and are thus unlikely to be properly loaded carboxysomes. Such non-fluorescent objects also display lower normalized scatter levels on average (Fig. S7), and they are excluded from further analysis. We also see some events with an anomalously high scatter and fluorescence such as event ii which may be attributed to aggregates. Interestingly, the event iii ends not by both signals going to background levels, but to a higher level on both indicating that the trapped object is replaced by a larger scattering carboxysome, highlighting how the ISABEL trap can only trap one object at a time. Many single carboxysome events like iv can now be analyzed.

The distribution of measured normalized scatter level for the trapped carboxysomes is shown in Figure 4e. The normalized scatter peaks at ~2.2 ×10^−4^, with a long tail to higher values. We can further visualize this distribution by looking at the density-colored scatter plot of fluorescence brightness vs normalized scatter for the same levels, as in Figure 4f. An extended distribution of scatter values covers one order of magnitude, with the fluorescence brightness covering nearly three orders of magnitude. The fluorescence brightness generally increases with normalized scatter, representing increased numbers of sfGFPs and thus increased core loading. This shape can be characterized by fitting a power law to the main density of trapping events (enclosed in the white dashed line in Fig. 4f, set as local density threshold). The power law fit in log space yields an exponent of 1.67, which deviates significantly from the linear relationship seen in the fluorescent polystyrene nanospheres. This unusual behavior arises from the complexity of the trapped object and will now be explored by modeling and additional measurements, in addition to extracting the mass of the entire carboxysome at the same time.

### Calculating carboxysome masses

The scattering cross-sections measured for the carboxysomes can be related to the masses of these objects using simple models for the scattering geometry and object parameters. With Mie theory, we calculate both the mass and the light scattering for carboxysomes assuming that they are spherical core-shell objects (Figure 5a), characterized by a total radius *r*_*total*_, a shell thickness *t*, and the densities of the core and shell, *ρ*_*core*_ and *ρ*_*shell*_, respectively. Here, the shell is assumed to comprise the hexameric shell proteins CsoS1A, B, and C (with minimal contribution from pentamers CsoS4A and B), and everything interior to the shell is assumed uniform and thus is averaged together into one core material. The refractive index increase over the refractive index of water is proportional to the additional density of proteins in that volume, represented by the total mass density *ρ* of proteins in *g*/*mL* as 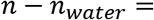 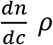 where 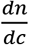 is the refractive index increment for proteinaceous material (see Methods). We use the average values *ρ*_*shell*_ = 0.96 *g*/*mL* and t = 3.5 nm for the shell density and thickness from crystal structures of the shell protein CsoS1A, assuming uniform thickness and shell packing across carboxysomes (Note S4), thus leaving the two free parameters *ρ_core_* and *r_total_* to characterize each carboxysome. With all four parameters, we can calculate the total mass of each carboxysome treated as a core-shell particle by multiplying the density and volume for both core and shell and adding the resulting core and shell masses.

The total carboxysome mass for relevant ranges of the total radius and core density can be calculated in MDa from the model (Figure 5b). The mass increases both with increasing *r*_total_ and with increasing *ρ*_core_. The scattering cross-section as a function of *r*_total_ and *ρ*_core_ can be calculated with Mie theory, and a contour of constant scattering cross section at a representative value of 2 nm^2^ is overlaid (black line in Figure 5b). A given value of scattering cross-section, as measured for an individual carboxysome in this experiment, can be achieved by increasing radius while decreasing density or vice-versa. Note that due to the size of the carboxysomes relative to the wavelength of light in water, *λ*_*water*_ = *λ*/*n*_*water*_ (*r*/*λ*_*water*_ ranges from 0.059 to 0.14), and thus the contribution of higher order terms in the Mie scattering calculation, the line of constant scattering cross-section is not also a line of constant mass. Put another way, these effects cause the mass to not scale exactly as 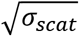. The mass along this constant cross-section contour is plotted vs radius in Figure 5c. For the range of densities and radii explored here, the mass corresponding to *σ*_*scat*_ = 2 nm^2^ first decreases slightly and then increases from 130 MDa to 160 MDa as the radius grows from 35 to 85 nm. For carboxysomes of a known radius and scattering cross-section, the mass is thus uniquely known.

In our experiment we do not directly measure radius, but we are able to measure a distribution of effective radii from cryo-TEM. Therefore, for a carboxysome with a measured scattering cross-section and a measured distribution of possible radii, below we will estimate a mass distribution by sampling from a mass curve analogous to Figure 5c. For a given carboxysome, we estimate its mass as the mean of this distribution.

The distribution of measured single-carboxysome scattering cross-section, calculated by applying Eqn (4) to the normalized scatter values in Figure 4e is shown in Figure 5d. This distribution is peaked roughly at 1.6 nm^2^ but extends past 15 nm^2^. The distribution of radii for circles of the equivalent carboxysome area measured by cryo-TEM for the same carboxysome sample is shown in Figure 5e and has a mean of 56 nm, and a standard deviation of 9 nm. We now use both these distributions to calculate the distribution of estimated mass values by resampling many values of the scattering cross-section from the experimental distribution (Figure 5d) and associating them with random values of the radius sampled from the distribution in Figure 5e. The calculated mass values are plotted against the associated scattering cross-sections in Figure 5f, with the radius of each calculated mass represented by the color. The major trend here is that the mass increases with increasing cross-section roughly as 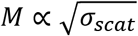, with the radius imparting a secondary effect. Not shown on the plot are some combinations of low scattering cross-section and large radius that are unphysical requiring a negative core density (<2%). The total distribution of deduced carboxysome masses from our measured cross-sections is shown in Figure 5g. The distribution is peaked at 130 MDa. We note that this distribution does not change substantially with changes in assumed shell thickness and shell density (Fig S8). To assess the uncertainty in calculated mass for a specific measured cross-section, we show the possible mass values for a 0.2 nm^2^ window around a scattering cross-section *σ*_*scat*_ = 2 nm^2^ (Figure 5g inset). This single value of scattering cross-section gives a mass distribution with mean 134 MDa and standard deviation of 5 MDa, demonstrating that even without knowing the precise radius of a given particle, the mass can be estimated to within 10%. Thus, we can measure the mass of a single carboxysome with minimal assumptions about its particular radius due to the weak dependence on radius in the relationship between mass and scattering cross-section.

### Modeling the fluorescence vs scatter relationship for internally-labeled carboxysomes

Finally, the simultaneous measurement of normalized scatter and fluorescence brightness of internally labeled carboxysomes provides additional information about the core loading of these carboxysomes. The assumption that the fluorescence from sfGFP fusion constructs targeted to the interior is proportional to the protein content of the carboxysome core, and thus the core mass in a core-shell model, further constrains the possible parameters describing the carboxysome ensemble. As we show below, these fluorescence measurements restrict the range of possible radii that could be associated with a scattering cross-section measurement.

There are two parameters we wish to explore: the radius and core density, constrained by our measurements of scattering cross-section. The first column of Figure 6 labeled a,b,c shows the case of no correlation between radius and scattering cross-section. Figure 6a shows the case of randomly picking a cross-section from Figure 5d and randomly picking a radius from Figure 5e as was done for the calculation of Figure 5f (see Note S5, Fig S9 for more details). The contours of constant probability show that the radius is approximately independent of scattering cross-section, however, for each point in this plot, the core density necessarily changes. We mark on the plot three cases and find the median radius can be extracted for 10% intervals of scattering cross-section (1.5 nm^2^, 3 nm^2^, and 6 nm^2^ marked i, ii, and iii, respectively). The calculated values for the core density matching the radius and scattering cross-section for these carboxysomes are shown in Figure 6b. The density increases from i to iii to account for increasing scattering cross-section as the radius remains constant. The core masses calculated for all the sampled carboxysomes are plotted in Figure 6c *vs* the normalized scatter calculated by using Eqn (4). (In this entire argument, the core mass is assumed to be a proxy for the fluorescence brightness.) The shape of this distribution can be characterized by extracting a power law analogous to Figure 4f, yielding in this case an exponent of 2.0. Note that as we proceed from i to ii and then to iii, the core mass increases as the core density increases, whereas the mass of the shell remains constant. The normalized scatter is more closely related to the total mass (approximately linear scaling), and thus since the shell mass is not changing, the core mass has to increase faster than the total mass which yields an exponent > 1 between core mass and normalized scattering signal.

**Figure 6.**
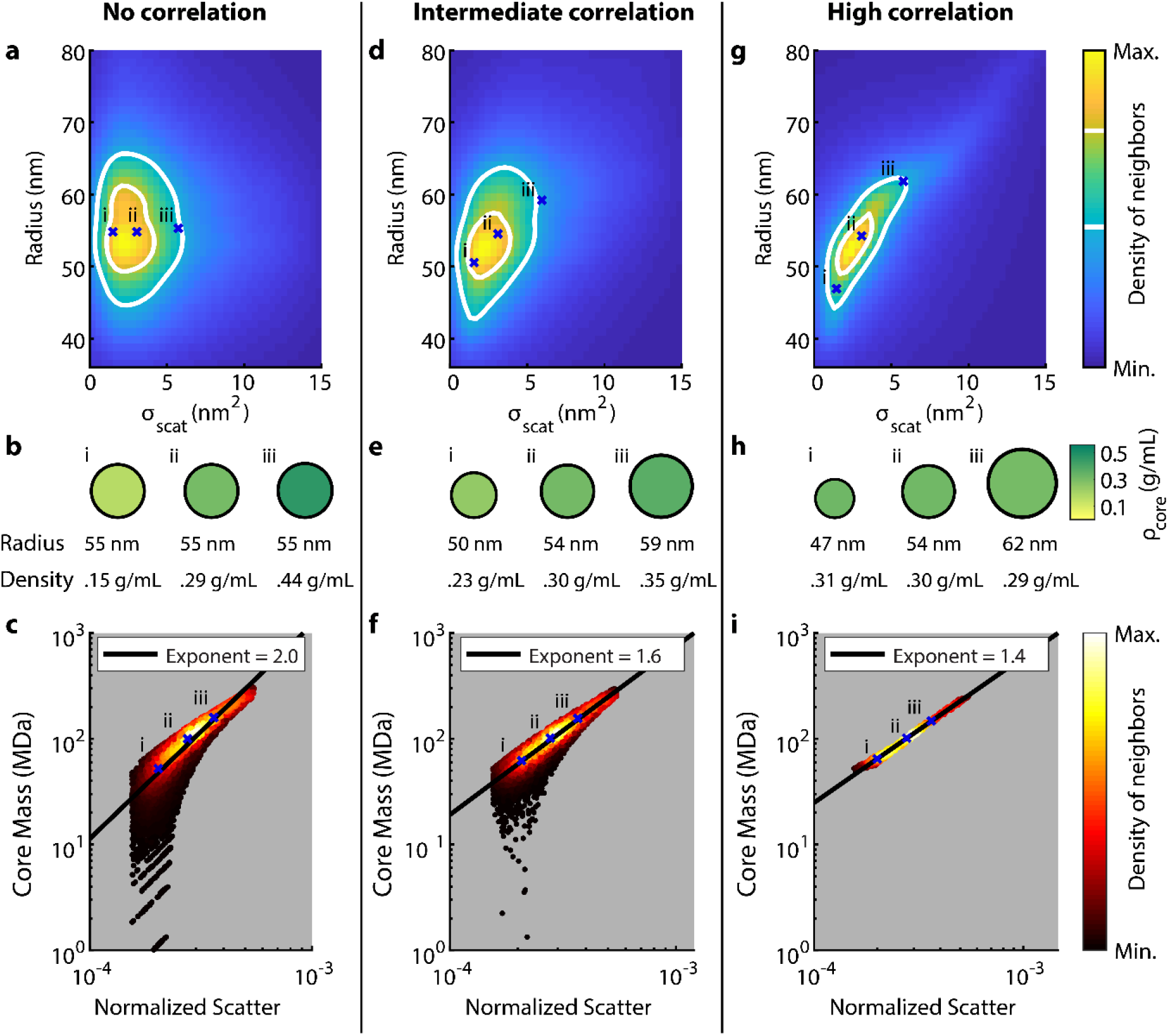
Simulating the fluorescence vs normalized scatter scaling for carboxysomes. (a,d,g) A density map for correlated values of radius and scattering cross-section generated from the measured univariate distributions with a correlation value of (a) 0, (d) 0.6, or (g) 0.99 by the method of copulas. Contours at 45% and 75% of max density are shown. The median diameters at 1.5, 3, and 6 nm^2^ are marked with a cross and labeled i, ii, and iii. (b,e,h) Carboxysomes with parameters marked in the above panels for each value of correlation. The total radius and core density calculated are written below. (c,f,i) The distribution of calculated core mass vs measured normalized scatter for model carboxysomes drawn from the given correlated distributions, with power-law fits overlaid. Carboxysomes with the parameters in a-c are marked with an ‘x’. The exponent in Fig. 6f approximately matches the measured exponent for the fluorescence vs normalized scatter distribution (Fig 4f) for an intermediate correlation value of 0.6.

The other extreme assumption would be that increased scattering cross-section is always associated with an increase in radius (third column of Figure 6 labeled g,h,i). This is enforced by closely matching the percentiles of the two variables when drawing values from the distributions (Note S5, Fig S9). Assuming a correlation of 0.99 for generating correlated radius and scattering cross-section distributions, the radius increases with increasing scattering cross-section with a narrow spread (Figure 6g). Carboxysomes with median radius are marked as before, and now show increasing radius from 47 to 54 and then 62 nm, while the core density remains nearly constant at ~0.3 g/mL (Figure 6h). Calculating the core mass and normalized scatter for each carboxysome (Figure 6i) now gives a very narrow distribution around a power law trend, with an exponent of 1.4. This decreased exponent reflects the fact that the increased normalized scatter is now caused both by increases in the mass of the fluorescent core and the mass of the non-fluorescent shell (as the radius increases). This exponent, however, is too small to explain the measured exponent for the fluorescence vs normalized scatter distribution in Figure 4f.

We can find an intermediate value for correlation used to generate the radius and scattering cross-section relationship in order to approach the exponent measured for the given carboxysome sample, in this case 0.6 (middle column Figure 6d,e,f). The distribution of radii and scattering cross-sections (Figure 6d) shows contours of constant probability density forming ellipses with slightly inclined axes. The median carboxysomes identified as before show increases in both radius and core density with scattering cross-section (Figure 6e). The distribution of core mass *vs* normalized scatter (Figure 6f) now better matches the distribution experimentally measured in Figure 4f, with an exponent of 1.6, intermediate between the uncorrelated and fully correlated cases. This suggests that increases in both the radii and core densities of the carboxysomes available contribute to an increase in scattering cross-section and mass. However, we note that while the radius and the core density are themselves positively correlated with increasing scattering cross-section, the radius is still anti-correlated with core density (Fig. S10). As well, we find the distribution of core masses for a single normalized scatter value in Figure 6f is still narrower than the distribution of fluorescence brightness for the same normalized scatter in Figure 4f (compared in Fig. S11), suggesting additional heterogeneity associated with these distributions. Nevertheless, by using the trendline of fluorescence vs scatter to constrain the unknowns, we can refine the distribution of carboxysome radii and narrow the distribution of core parameters, learning more about the distribution of mass within a carboxysome.

## Discussion and Conclusion

Taking advantage of the relation between interferometric scattering signal and protein mass, we have extended the ISABEL trap to include mass measurements of single trapped objects. To do this, we first demonstrated the single-particle scattering cross-section measurement capabilities by calibrating the scattering with well-defined single-component beads. We validated these calibrations by inferring the diameters of polystyrene nanospheres, with the measured diameters lying within 2% of the values measured by cryo-TEM. By combining the measurements of scattering cross-section with parameters of the carboxysome geometry known from EM and structural studies, we can also calculate the masses of single carboxysomes with ~10% uncertainty.

Our simultaneous fluorescence measurements of GFPs targeted to the carboxysome cargo can exclude non-fluorescent objects from study, and the shape of the fluorescence vs scatter distribution can also be understood in a core-shell model with a simple assumption about the relation between increasing scatter and radius. The comparison of our measured masses and mass distributions to previous publications is complicated by the sensitive dependence of the carboxysome loading on the preparation conditions. For example, comparing the *H. neapolitanus* carboxysomes measured with STEM^12^ and cryo-ET^13^ to our carboxysomes grown in *E. coli*, we see a 10% increase in the carboxysome diameter, while *H. neapolitanus* native and *E. coli* expressed carboxysomes in a work characterizing stoichiometry^47^ are 10% larger still, suggesting that the assembly process is sensitive to growth conditions. Nevertheless, our measured mass values of 100 MDa to 250 MDa are in the same range as 160 MDa to 260 MDa reported in the work by Schmid *et al*.,^12^ while the larger carboxysomes of Sun *et al.*^47^ are on average 340 MDa. The best estimate of our core mass is also in the range 50 MDa to 150 MDa (Fig. S12) with core densities around 0.25 g/mL (Fig. S13). Note that our results are somewhat dependent on the choice of shell parameters (Fig S14, S15 and S16). In comparison, the population averaged core mass in Sun *et al.* was 273-286 MDa, which is proportionally greater as a fraction of the total mass (~82% compared to ~65% in this work). These results can be additionally compared to cryo-ET results from Metskas *et al.*^13^ if we assume that the rubisco proteins measured are a proxy for the total core mass, who report rubisco concentrations of 300 μM to 900 μM, with total numbers 150 to 400. These can be converted to rubisco mass concentrations and totals of 0.16 g/mL to 0.49 g/mL and 81 MDa to 216 MDa, respectively. These comparisons show the clearest points of difference for interferometric scattering over cryo-ET – where cryo-ET can provide exquisite structural detail for objects that can be identified, the subsequent mass calculation will miss contributions from proteins that are not clearly identified in cryo-ET, such as the intrinsically disordered scaffolding protein CsoS2.^48^

The largest current limitation on mass and composition measurements in the ISABEL trap is an accurate simultaneous determination of the total radius. A simultaneous measurement of object radius can be derived for example by analogy to hydrodynamic radii estimated from diffusion measurements in the ABEL trap^49^ or by using single-particle tracking (without trapping^22^). Such data in addition to measurement of scattering cross-section would alleviate the need for assuming the fluorescence to be proportional to the total cargo mass, and could enable the study of the variability in cargo composition. Increasing the incident power would also decrease the size of the smallest objects that could be studied. Compared to other light scattering methods, the ISABEL trap has the benefit of simultaneous spectroscopy capabilities over extended time periods (multiple seconds) as a consequence both of the near-infrared scattering wavelength and the extended observation without tethering. We can already measure the redox behavior inside carboxysomes in the ISABEL trap^31^, and have measured detailed photophysical and kinetic parameters inferred from fluorescence in previous ABEL trap studies.^27^, ^29^ These measurement capabilities could be combined to apply to many types of biological microcompartments with an enclosing shell, loaded cargo, and sequestered internal chemistry. The combination of scattering detection and particular fluorescence labels could enable the characterization of all such objects produced in a biological system, uncovering yet more of their hidden heterogeneity.

## Supporting Information Description

See supplementary material for notes on the optical layout, analysis details, calibration of scattering cross-section, calculating carboxysome shell thickness and density, and generating correlation distributions of scattering cross-section and radius. A supplementary table summarizes the nanoparticle diameters. Supplementary figures show distribution of diameters measured for the 0.1 μm Fluospheres and 135 nm polystyrene nanospheres, the distribution of scattering cross-sections used for calibration, the optical layout schematic, the fractional error in normalized scatter measurements vs time, the effect of calculation choice on measured normalized scatter levels, the carboxysome levels excluded from analysis, the effects of changing shell parameters on the estimated mass, the schematic for generating correlation distributions of scattering cross-section and radius, the distribution of core densities vs radius, the comparison of the width of fluorescence and core mass distributions, the distribution of core masses, the histogram of core densities for best estimate, the effects of changing shell parameters on the required correlation, the effects of changing shell parameters on the estimated core mass, the effects of changing shell parameters on the estimated core density, and the comparison of estimated masses across technical replicates.

## Acknowledgments

This work was supported in part by the U. S. Department of Energy, Office of Science, Office of Basic Energy Sciences, Chemical Sciences, Geosciences, & Biosciences Division, Physical Biosciences Program, under Award Numbers DE-FG02-07ER15892 (W.E.M.) and DE-SC0016240 (D.F.S.). D.D.P. was supported by a Stanford Graduate Fellowship.

## Data Availability

The data that support the findings of this study are available from the corresponding author upon request.

