## Supporting Information for "Exploring masses and internal mass distributions of single carboxysomes in free solution using fluorescence and interferometric scattering in an anti-Brownian trap"

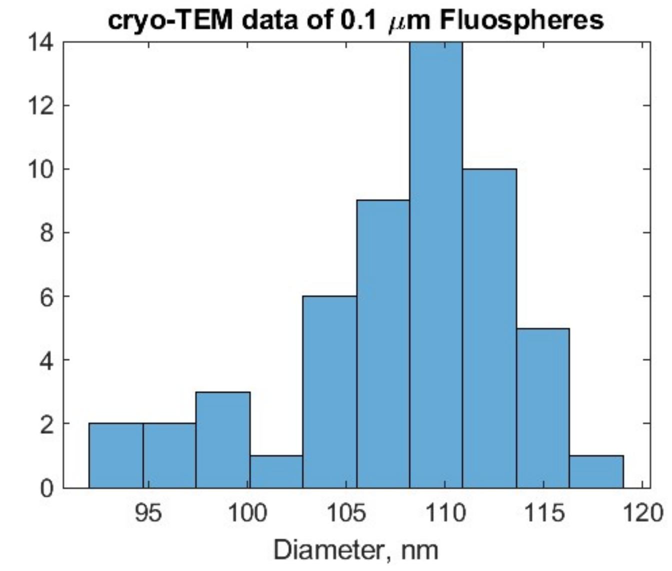

*Figure S1* Distribution of diameters measured with cryo-TEM for the Fluospheres with nominal diameter 0.1 μm. The measured mean and standard deviation were 107.9 nm and 5.7 nm for 57 nanospheres.

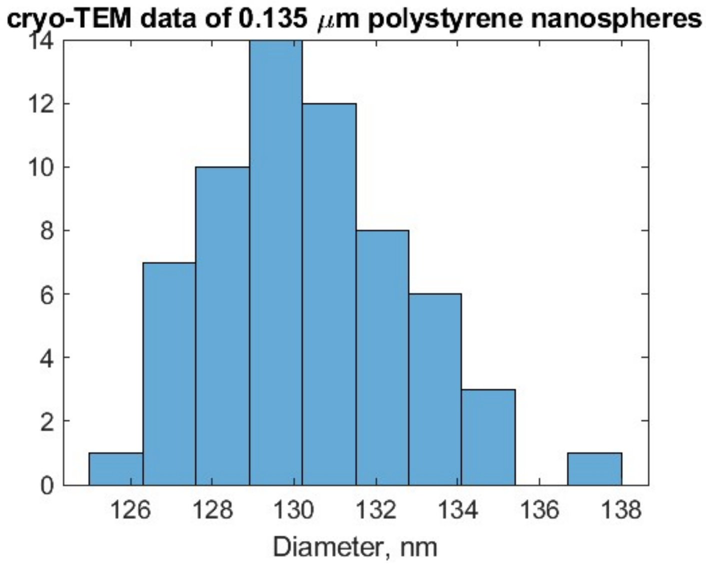

*Figure S2* Distribution of diameters measured with cryo-TEM for the polystyrene nanospheres with nominal diameter 0.135 μm. The measured mean and standard deviation were 130.4 nm and 2.4 nm for 62 nanospheres.

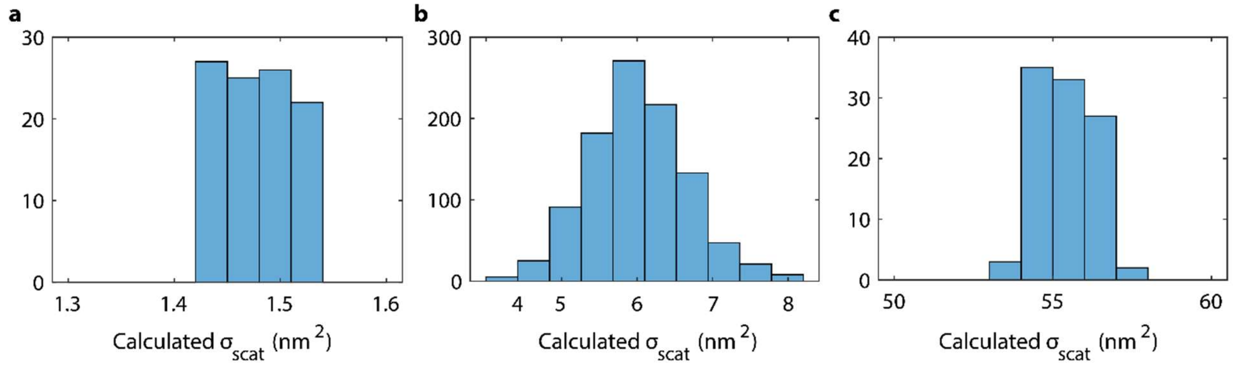

*Figure S3* Scattering cross-sections calculated with Mie theory for the nanosphere samples used for calibration. (a) Gold nanospheres of nominal diameter 30 nm: $\sigma_{scat} = 1.48 \pm 0.3$ nm$^2$ (s.d.). (b) Polystyrene nanospheres of nominal diameter 80 nm: $\sigma_{scat} = 6.0 \pm 0.6$ nm$^2$. (c) Gold nanospheres of nominal diameter 50 nm: $\sigma_{scat} = 55.4 \pm 0.9$ nm$^2$.

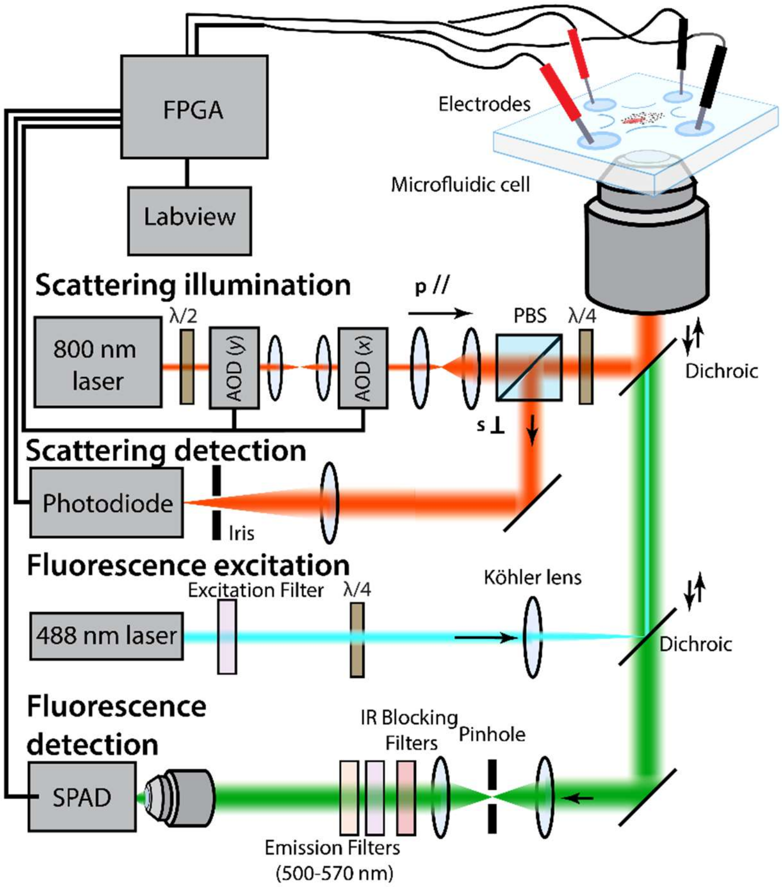

*Figure S4* Schematic of the optical paths of scatter and fluorescence beams, described in more detail in Note S1. The scatter illumination beam is deflected by two acousto-optic deflectors (AODs) controlled by the FPGA; it is linearly polarized at the polarizing beam splitter (PBS), then circularly polarized with a quarter-wave plate (λ/4). The back-reflected beam is orthogonally polarized after the quarter-wave plate and exits the other port of the PBS, and is ultimately focused for detection on a photodiode. Position is monitored and feedback voltages are calculated on the FPGA, then applied to the solution with platinum electrodes. Fluorescence emission spanning 500-570nm is collected on a single photon avalanche photodiode (SPAD) after spatial filtering with a 100 µm pinhole. Detected photons represented by TTL pulses from the SPAD are recorded and time tagged on the FPGA. AOD: acousto-optic deflectors, PBS: polarizing beam splitter, DC: dichroic beamsplitter, λ/2: half-wave plate.

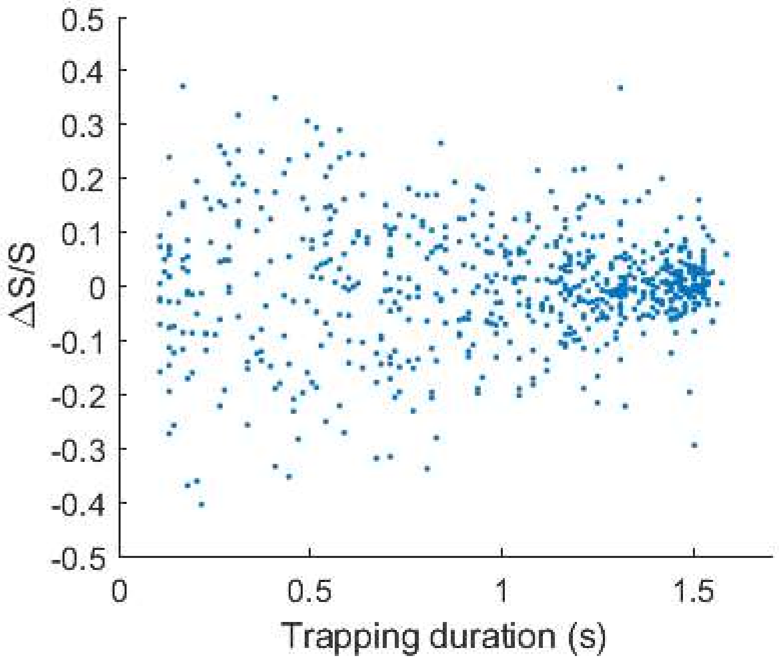

*Figure S5* Fractional error in normalized scatter vs duration of trapping event. The standard deviation of all the fractional errors plotted here is 0.13. See Note S2 for further details.

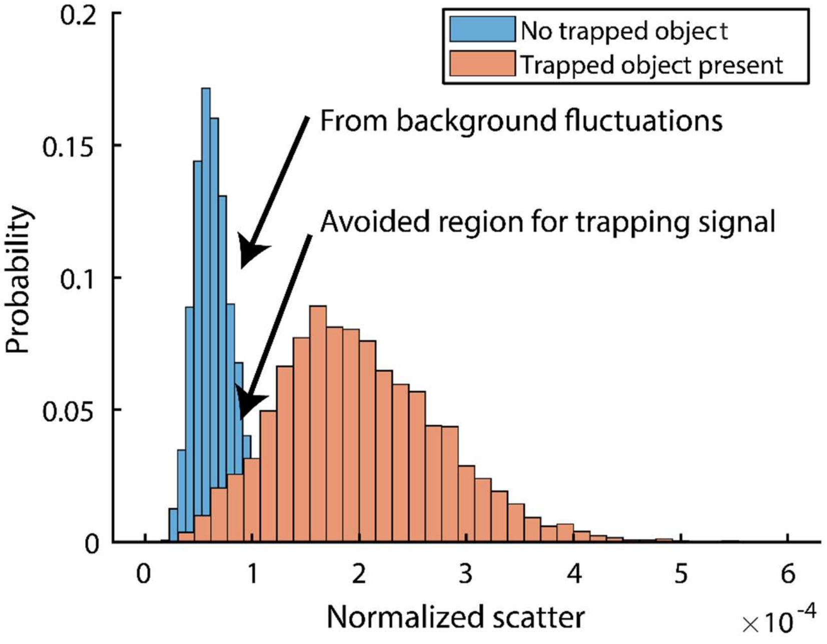

*Figure S6* Histograms of the frame-by-frame normalized scatter for the ISABEL trap with no object trapped vs frame-by-frame normalized scatter for a single, small trapped object (30 nm gold bead). The local maximum (center 3x3 beam positions for each frame) is unlikely to dip below the maximum of the background fluctuations, and thus systematically increases normalized scatter averages contributing to a level. See Note S3 for further details.

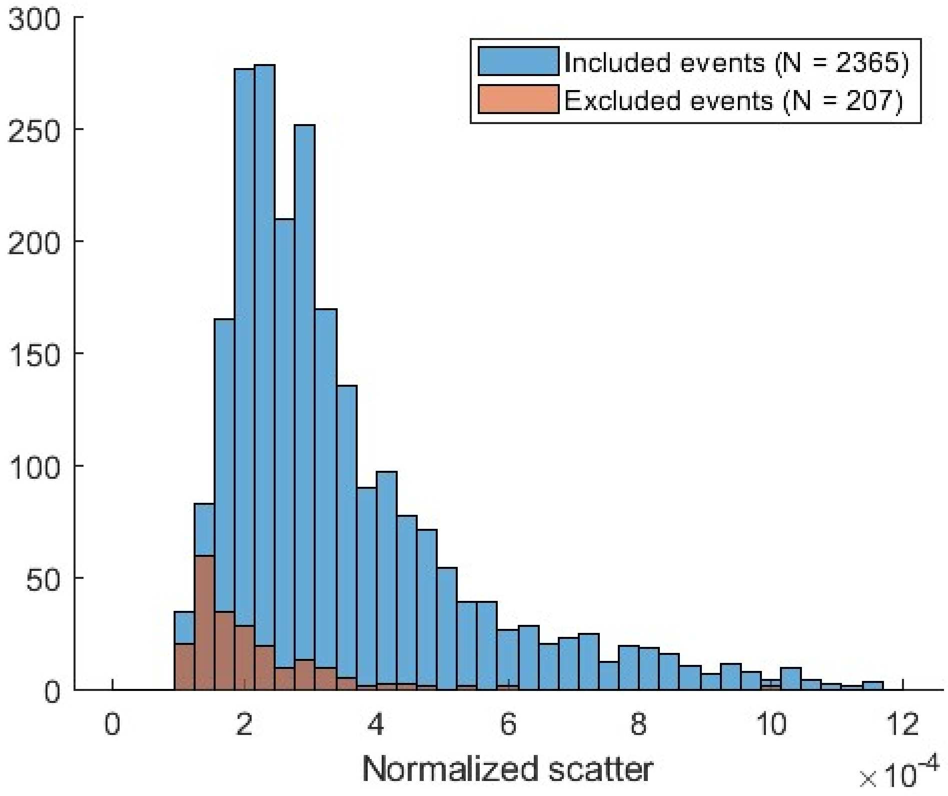

*Figure S7* Histogram of normalized scatter values during carboxysome trapping for events included and excluded from further analysis based on non-zero fluorescence brightness after background subtraction. The included events are plotted in Figure 4e. Excluded events have a lower normalized scatter on average.

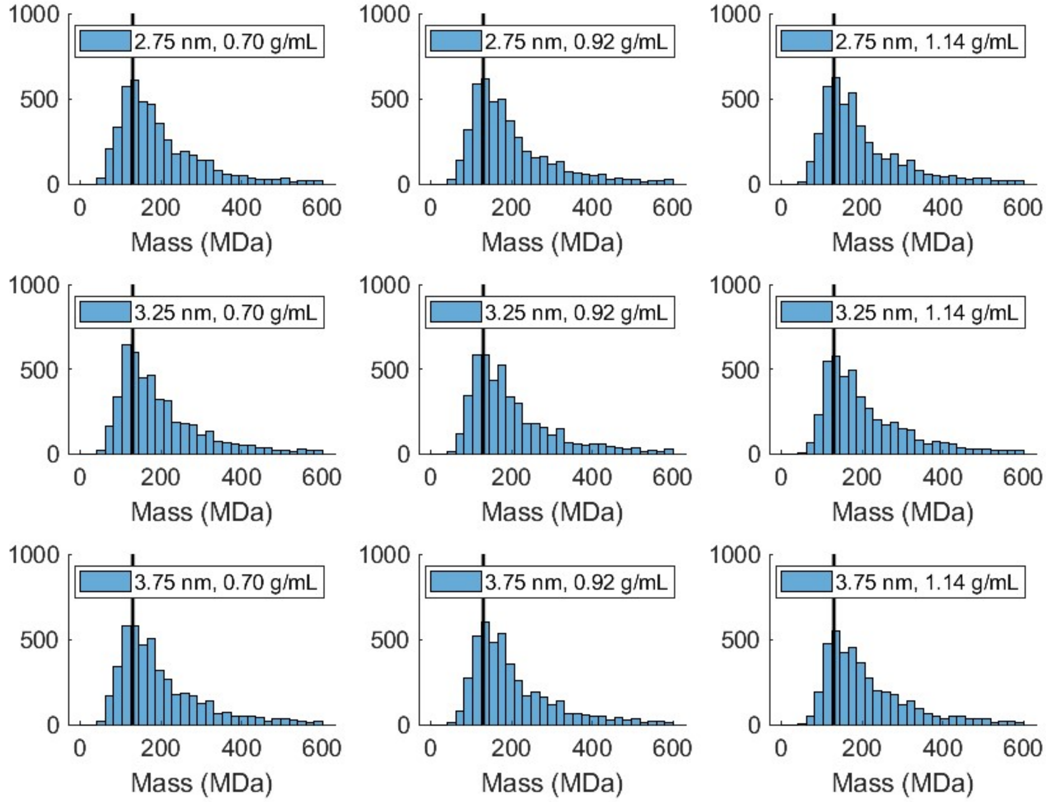

*Figure S8* The mass histograms for different values of shell thickness and shell densities, calculated in the same way as Figure 5g. A line at 130 MDa is drawn to aid the eye. There is no significant shift between the total mass values.

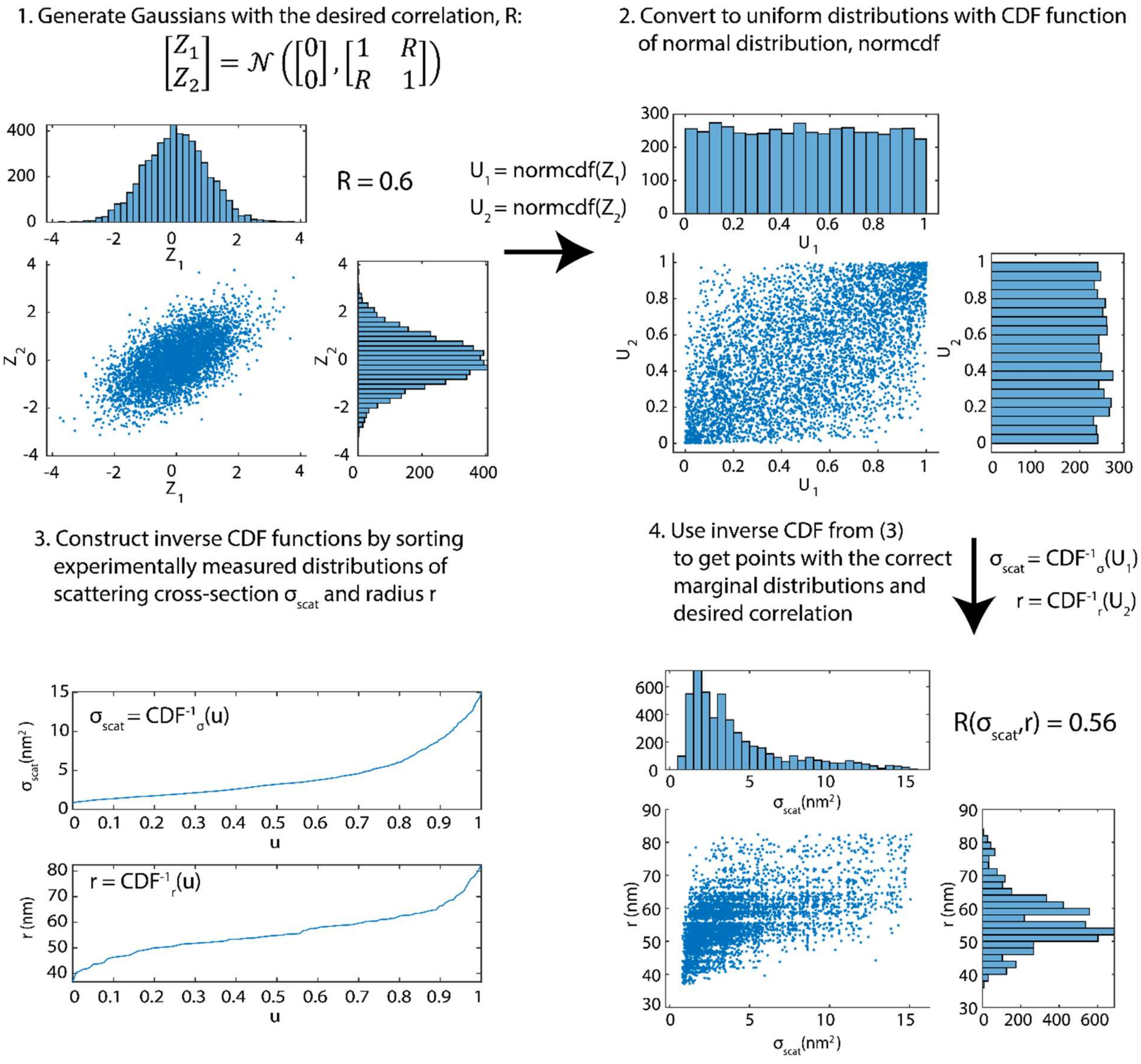

*Figure S9* Generating correlated values of scattering cross-section and radius matching the measured distributions by the method of copulas.[1] This method relies on generating two correlated Gaussian random variables, and then transforming each of the two random variables into the desired distributions. 1) 5000 samples of a pair of correlated Gaussian random variables ($Z_1$, $Z_2$) are generated with a correlation R = 0.6. 2) Applying the CDF function for a normal Gaussian variable independently to each of the two gives uniformly distributed random variables $U_1$ and $U_2$, that are correlated. 3) Determining a one-to-one map from the unit interval [0, 1] to the experimentally measured $\sigma_{scat}$ and $r$ distributions requires the inverse CDF functions of these variables $CDF_{\sigma_{scat}}^{-1}$ and $CDF_r^{-1}$ , which can be generated by interpolating the sorted list of measurements vs a normalized index u. 4) Applying the experimental inverse CDF to the correlated uniform variables ($U_1$, $U_2$) determined in (2) gives pairs of sampled $\sigma_{scat}$'s and $r$'s that match the experimental distributions, but have an underlying correlation. See Note S4 for further details.

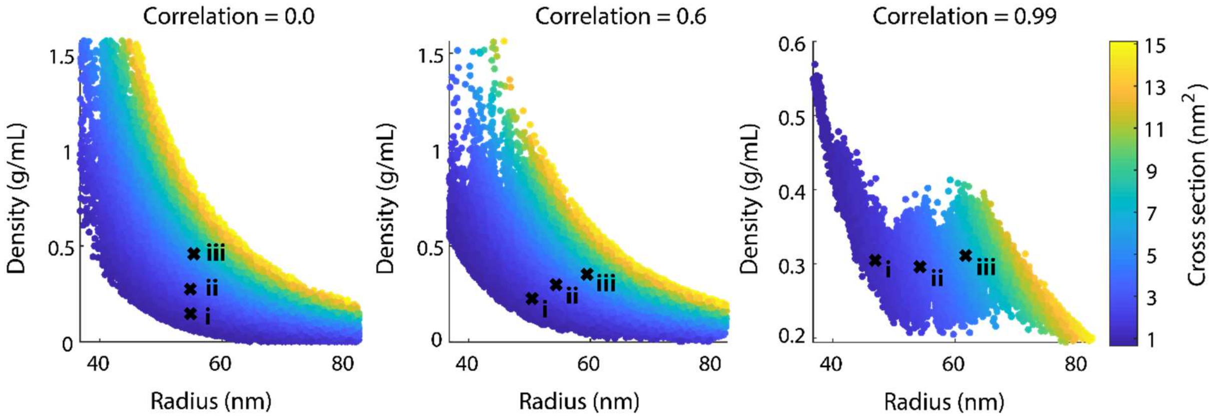

*Figure S10* Distribution of core density vs radius for different correlation values. The datapoints are the same as displayed in Figure 6a,d,g, and the points (i, ii, iii) are also marked here.

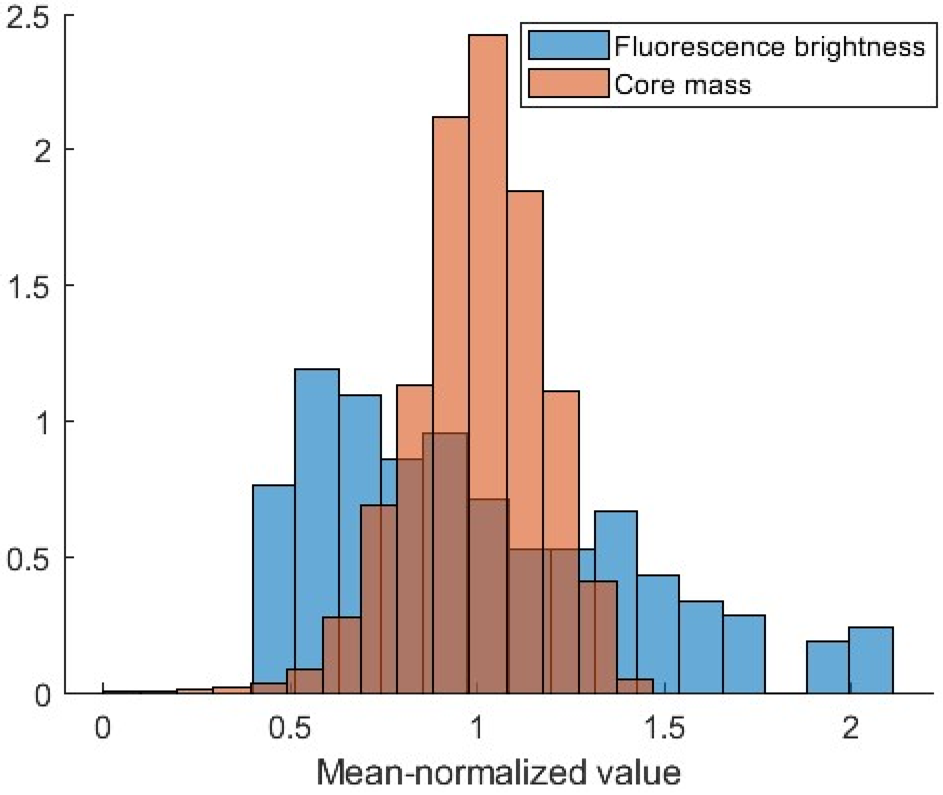

*Figure S11* Comparing widths of the normalized fluorescence brightness $F/\langle F\rangle$ measured for the carboxysomes in a 10% window around normalized scatter value of $\mathcal{S} = 2.2 \times 10^{-4}$, to the normalized core mass $m_{core}/\langle m_{core}\rangle$ calculated for the model carboxysomes in Figure 6f with correlation R = 0.6, also in a 10% window around $\mathcal{S} = 2.2 \times 10^{-4}$. Core masses show a much narrower distribution than the fluorescence brightnesses. The additional width of the fluorescence distribution may reflect loading heterogeneities of the sfGFPs compared to the main core cargo.

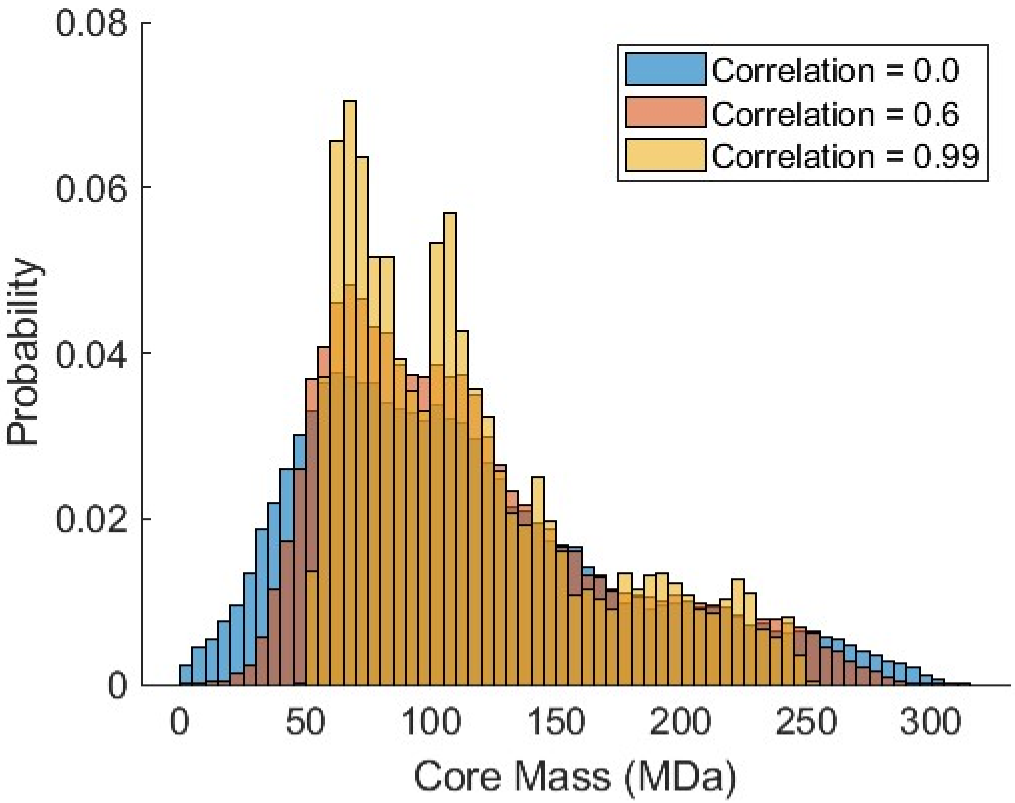

*Figure S12* Histograms of core masses calculated for the experimentally measured carboxysomes generated with correlation values 0.0, 0.6, and 0.99 between radius and scattering, from Figures 6c,f and i.

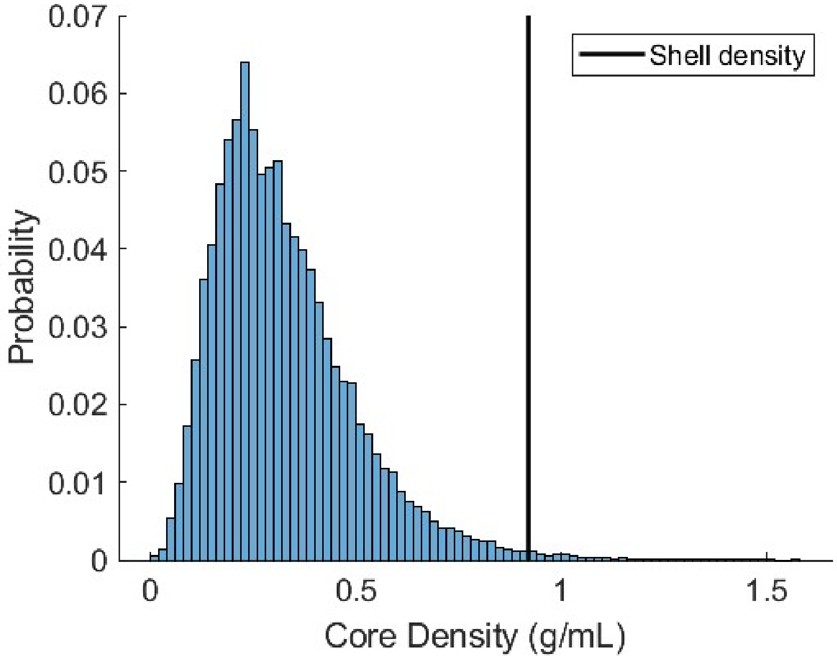

*Figure S13* Histogram of core densities for carboxysomes calculated in Figure 6f, with correlation 0.6 between radius and scattering cross-section. Vertical line is at the value of shell density used.

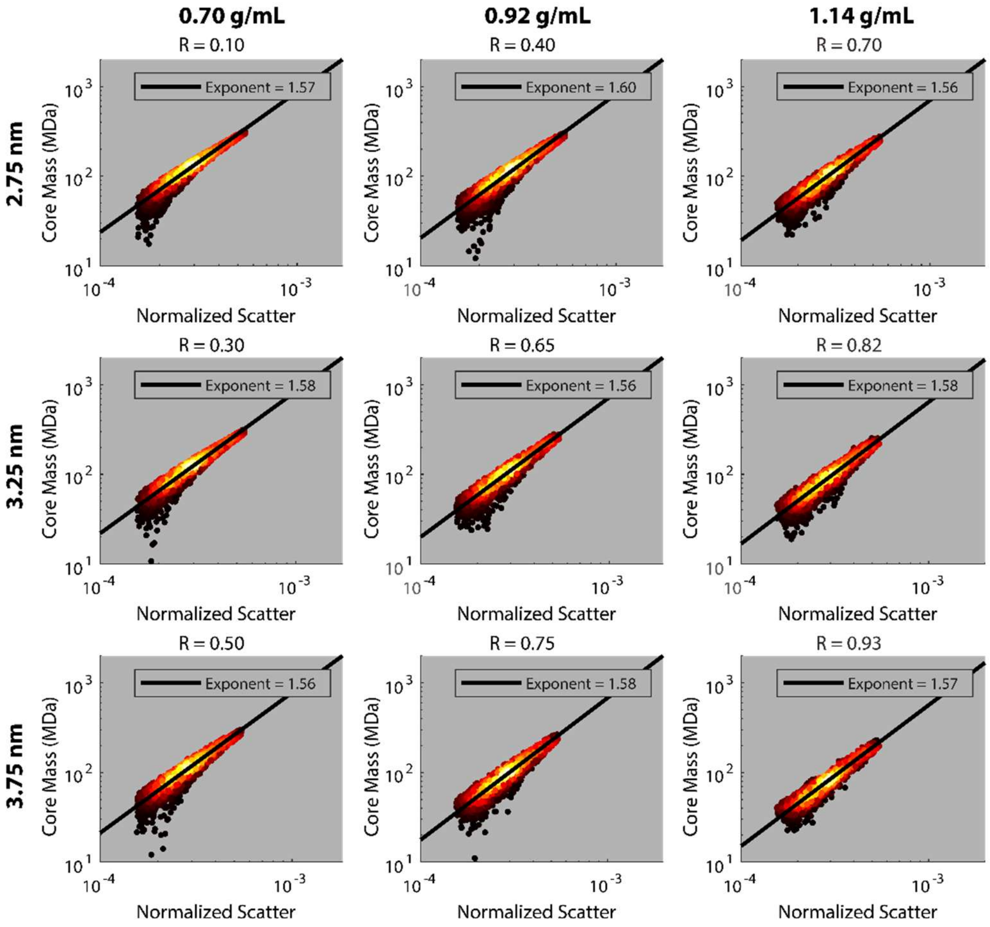

*Figure S14* Effect of changing shell parameter assumptions. The different assumptions shown require changing the correlation R used to generate core mass *vs* normalized scatter plots matching the experimental trends. As the shell mass increases by increasing either the shell thickness (top to bottom) or the shell density (left to right) around the value used in the main text (center), the correlation value R must also be increased to roughly match the experimental exponent for fluorescence vs normalized scatter, 1.57.

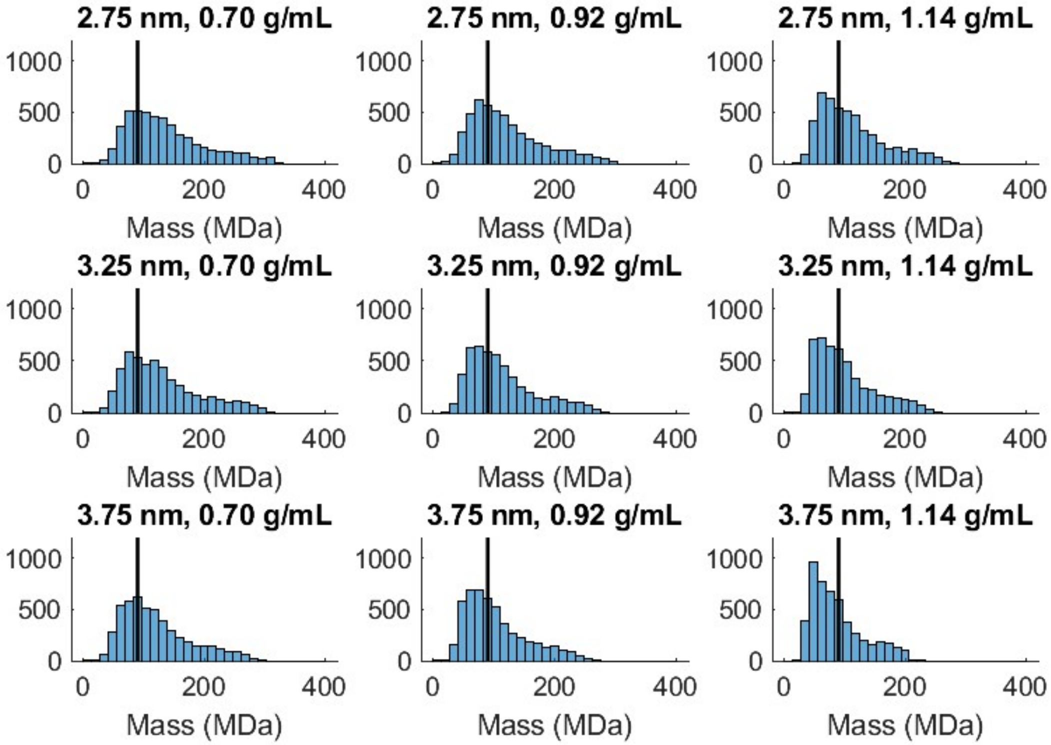

*Figure S15* Estimated core mass distributions as the shell parameters are varied. The distribution of core masses for the shell parameters and correlation values in Figure S14 above is plotted. The line is drawn to aid the eye at 90 MDa on each plot. When the trendline exponent is matched to experimental data, the core mass distributions shift to lower masses as the shell thickness or density is increased.

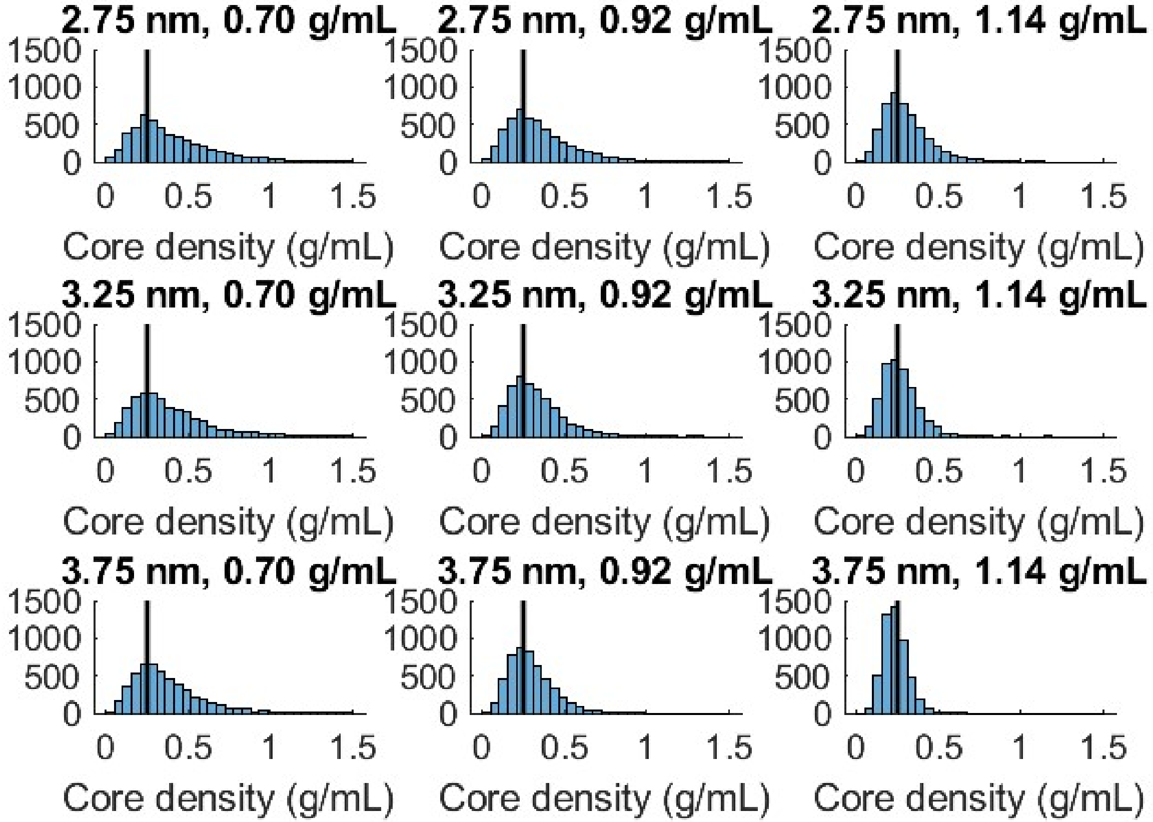

*Figure S16* Estimated core densities as shell parameters are varied. The distribution of core densities for the shell parameters and correlation values in Figure S14 above is plotted. The line is drawn to aid the eye at 0.25 g/mL on each plot. If the experimental trendline exponent is matched, the core density distribution remains at the same peak value, but these distributions narrow with increasing shell thickness and density.

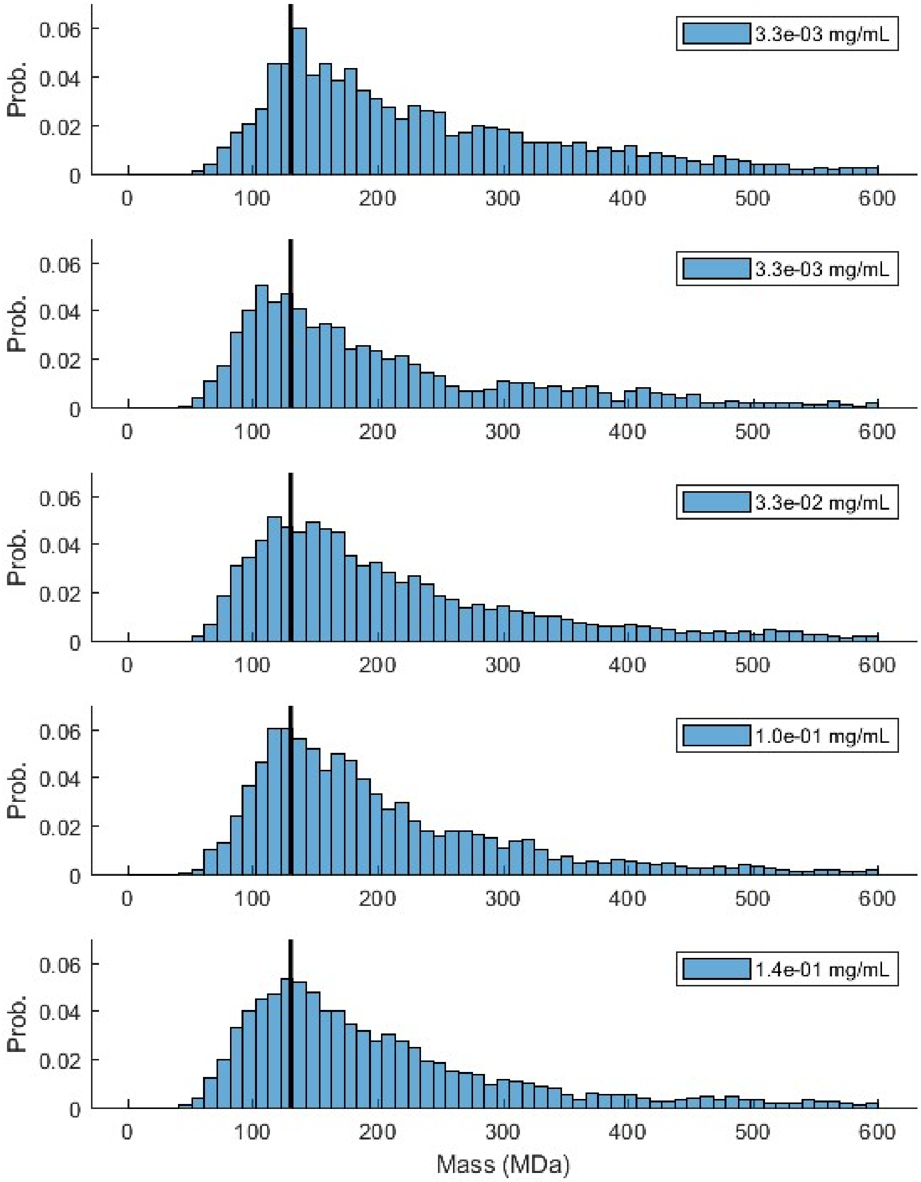

*Figure S17* Comparing five different datasets measuring carboxysome masses. The samples were prepared identically as described in the text, with changes in the concentration of antibodies used: 3.3 µg/mL (N = 733), 3.3 µg/mL (N = 998), 33 µg/mL (N = 2458), 100 µg/mL (N = 3362), and 140 µg/mL (N = 2347, same dataset as Figure 5g). The vertical line is drawn at 130 MDa to aid the eye. Note that the total mass displays no discernable trend with increasing antibody concentration, suggesting that the antibodies do not contribute measurably to the mass.

*Table S1.* Summary of nanoparticle diameters. The cryo-TEM measured diameters for the 0.1 µm Fluospheres and 135 nm polystyrene nanospheres were scaled to match the mean diameter for the NIST-traceable nanospheres.

| **Sample** | **Manufacturer provided diameter: mean and std. dev.** | **Cryo-TEM measured diameter: mean and std. dev. (this work)** |
|---|---|---|
| 30 nm gold | 28.0 ± 0.9 nm | - |
| 50 nm gold | 51.0 ± 0.9 nm | - |
| 80 nm polystyrene | 81 ± 9.5 nm (NIST-traceable to 81 ± 1.5 nm) | 79.3 ± 6.1 nm (N = 53) |
| 0.1 µm Fluospheres | 0.1 ± 0.0063 µm | 107.9 ± 5.7 nm (N = 57) |
| 135 nm polystyrene | 0.135 µm (no std. dev quoted) | 130.4 ± 2.4 nm (N = 62) |

**Note S1: Optical layout**

The optical layout is a slight modification of the previously published ISABEL trap setup.[2] For completeness, we repeat most of the details here.

<u>Interferometric scattering channel:</u> Briefly, the NIR illumination is provided by a multimode 802 nm laser diode (Axcel M9-808-150-D5P) driven by a Thorlabs LDC 500 laser diode driver, housed in a Thorlabs TCLDM9 mount, and temperature stabilized by a Thorlabs TEC2000 thermo-electric cooler driver. The trapping is made possible when an additional sinusoidal voltage is applied to the diode through a bias tee in the diode mount as in Ref.[3]. This modulating signal was provided by a function generator (Agilent 33220A) at 2.5 MHz with $V_{pk-pk}$ = 2.00V, which corresponds to a modulation current of 40 mA peak-to-peak on top of the 120 mA DC drive current from the laser diode driver. The corresponding reduction in coherence removes fluctuations of the background, likely arising from mechanical drifts in the system which produce changes in interference between weak reflections. The beam is roughly circularized with

a 4:1 anamorphic prism pair and is then further spatially filtered with a single-mode fiber (ThorLabs P1-780A-FC-5).

The polarization after the single-mode fiber is controlled with a half-wave plate (HWP) which is adjusted daily. The beam is then appropriately magnified with a telescope and passed through a pair of AODs (AA Opto Electronics MT110-B50A1,5-IR, driven by DDSPA2X-D4125b-34) that are programmed by the FPGA to drive the beam in a predetermined scan pattern in the sample plane. The beam is moved to a new position in the Knight's tour pattern every 18.75 μs. A pair of lenses image the turning plane in the first AOD to the turning plane in the second AOD, with another pair of lenses mapping this plane onto the back focal plane of the objective, such that beam is deflected with no displacement in the back focal plane and thus translated with no turning in the sample plane after passing through the objective. The beam travels through a polarizing beam splitter (PBS) and a quarter wave-plate (QWP), and is reflected off a 775 short-pass dichroic (Chroma) on the way to the objective (Olympus UPlanApo Oil-Iris, oil-immersion, 100X, 1.35NA). The HWP is adjusted to maximize transmission through the PBS. The QWP is adjusted to maximize the back-reflection signal heading to the detector. This reflection from the sample plane is focused onto a large-area photodetector (Newport 2031). An adjustable iris close to the detector allows only light near the desired reflection from the quartz-water interface to pass to the detector, and a 90:10 beam splitter (not shown in the schematic) allows some of the light to be imaged on a Si CMOS camera (FLIR Blackfly S U3-19S4M) to check alignment, focus, and iris positioning. The photodiode is set at its medium gain setting for most experiments with a transimpedance gain of $10^5$ V/A and a bandwidth of 150 kHz. To allow the photodiode signal to settle and to reduce noise, starting 10 μs after the beam is moved, the photodiode signal is read by the FPGA board (NI RIO PCIe-7856R) eight times 1 μs apart and these values are averaged to define the detected power from the current beam position.

Fluorescence excitation and detection: The 488 nm excitation laser (Coherent Sapphire, 100 mW) is sent through an excitation filter (Chroma, 488/3) and circularized in its polarization by a quarter-wave plate. The laser power is adjusted with neutral density filters to give powers between ~1 and ~10 μW at the sample plane. A Köhler lens provides an excitation spot in the sample that covers the entire trapping region, with $1/e^2$ radius of 5.5 μm.

In the fluorescence collection path, a pinhole is used to restrict the collection area to cover the entire 3.6 μm diameter Knight's tour pattern. Blocking filters (808 nm notch reject, 785 nm short pass are provided to prevent the 800 nm from interfering with the fluorescence detection. Additional 488 nm long pass, 500 nm long pass, and 570 nm short pass filters process the final fluorescence signal before detection on the photon counting detector (Picoquant τ-SPAD).

**Note S2: Analysis Notes**

Identifying trapping event signals offline: The levels associated with a single trapping event are identified using a maximum-likelihood change-point algorithm with a Gaussian model[4] on the normalized scatter data, binned for every 20 frames (12 ms). Consecutive levels within 10% of each other are combined. Normalized scatter levels are identified as trapping events if the following three conditions are met: (1) it lasts longer than 0.1 s, (2) it ends within 150 ms of the feedback being turned off, and (3) the measured position has a frame-to-frame RMS value with respect to the trap center less than 1.2 μm. The second condition allows some time for objects to diffuse away when feedback is toggled off, but excludes objects stuck to the quartz surfaces. For the simultaneous fluorescence data, the fluorescence brightness level was calculated (during the level identified from the scattering signal) as the median of the 10ms-binned fluorescence brightness, and an empty-trap fluorescence background level was subtracted. The fluorescence background level for a trapping event was identified as the lowest fluorescence level of the two preceding and two succeeding empty-trap levels.

Estimating error in a trapping event: The normalized scatter shows large fluctuations from frame to frame, including changes in the relative phase term in the interferometric contrast (Eqn (4) in the main text). We estimate the fractional error in a trapping event with the difference in the normalized scatter $\mathcal{S}$ averaged over the first half and over the second half of the trapping levels identified for nominal 50 nm gold nanospheres, see Figure S2:

$$\frac{\Delta\mathcal{S}}{\mathcal{S}} = \frac{\langle\mathcal{S}(t_i < t < t_{i+1/2})\rangle - \langle\mathcal{S}(t_{i+1/2} < t < t_{i+1})\rangle}{\langle\mathcal{S}(t_i < t < t_{i+1})\rangle}$$

The distribution of fractional errors narrows for longer trapping durations, suggesting that longer measurement times continue to improve precision on measured scatter signal, likely due to more thorough sampling from the distribution of mode overlaps and phase factors. Since the fractional error is the difference between two noisy values, we expect it to be a factor of $\sqrt{2}$ larger than the precision of the normalized scatter measurement on half the level. Fig. S2 demonstrates that for trapping times greater than 1s, the fractional uncertainty in a normalized scatter level is less than ~10%.

Analyzing fluorescence brightness vs normalized scatter trends: The local densities of points were calculated in log-log space. The power-law fits to fluorescence brightness *vs* normalized scatter were determined with a linear fit in log-log space. The fit to the measured carboxysome data in Figure 4f applied a cutoff of 25% of max density, maximum normalized scatter $5.5 \times 10^{-4}$, minimum fluorescence brightness 30 photons/10ms, and maximum fluorescence brightness 2000 photons/10ms. The fits to core mass *vs* normalized scatter for simulated carboxysomes in Figure 6c,f,i were also performed on log-log axes, with no cutoffs. The results for fitting the exponents are summarized below:

| Sample ID | Best-fit exponent | 95% confidence intervals |
|---|---:|---:|
| 0.1 μm Fluospheres | 0.93 | (0.865, 0.995) |
| Carboxysomes (experimental) | 1.57 | (1.504, 1.635) |
| Carboxysomes (simulated, R = 0.0) | 2.038 | (2.029, 2.048) |
| Carboxysomes (simulated, R = 0.6) | 1.593 | (1.589, 1.598) |
| Carboxysomes (simulated, R = 0.99) | 1.377 | (1.376, 1.378) |

Given the confidence intervals on the exponent for the experimental data in Figure 4f, the exponents were analyzed to only two significant figures.

**Note S3: Calibrating scattering cross-section *vs* normalized scatter level**

Fitting a power law for calibration: Values of the normalized scatter and scattering cross-section for each of the three calibration samples (30 nm gold nanospheres, 80 nm polystyrene nanospheres, and 50 nm gold nanospheres) were drawn from Gaussian distributions in log space with the following parameters:

| **Sample ID:** | **Gold, 30 nm** | **Polystyrene, 80 nm** | **Gold, 50 nm** |
|---|---|---|---|
| $\log(\sigma_{scat})$ | 0.168 ± 0.012 | 0.778 ± 0.046 | 1.744 ± 0.0076 |
| $\log(\mathcal{S})$ | -3.704 ± 0.029 | -3.417 ± 0.087 | -3.024 ± 0.021 |

The best fit straight line was obtained from the three points, and the process was repeated to obtain 1000 samples of straight lines for $\log(\sigma_{scat}) = a\log(\mathcal{S}) + b$. Recasting the equation in the following form and calculating means and standard deviations for the 1000 samples we arrive at the calibration expression:

$$\sigma_{scat} = \left(\frac{\mathcal{S}}{1.71 \pm 0.04 \times 10^{-4}}\right)^{2.33 \pm 0.02} \text{nm}^2$$

Extracting normalized scatter values from trapping data, and the impact of this definition on the calibration curve: The method of determining the normalized scatter signal for a frame, by taking the maximum normalized scatter from the central 3 x 3 beam positions around the nominal trap center, systematically overestimates the average normalized scatter for smaller scatterers. This can be seen in Figure 2a, by comparing the signal from the first nanoparticle trapping event to the signal before it was trapped. The frame-by-frame signal never explores the full range between the max value (resulting from optimized mode overlap factor $M$ and phase factor $\cos\theta$) and zero, since the maximum of an absolute value cannot be zero. The minimum value explored by the frame-by-frame average is affected by the analysis procedure, and is greater than what would occur for pure background, the maximum of the absolute value of nine Gaussian random variables (Fig. S4). Thus, the average of the normalized scattering signal over a level is systematically increased for smaller objects. This is consistent with the fact that the calibration curve for scattering cross-section as a function of normalized scatter yields an exponent > 2.

**Note S4: Carboxysome shell thickness and density**

Estimating shell thickness and density from published crystal structures: The shell parameters were estimated by considering the density inside and thickness of a bounding box of a CsoS1A hexamer as follows. The CsoS1A monomer structure[5, 6] was analyzed in PyMol, converted to a hexamer by the symexp function in PyMol.[7] A bounding box for the hexamer was calculated,[8] with dimensions 7.21 nm × 7.38 nm × 3.22 nm. The shell thickness then can be estimated from the z-dimension of the box, 3.22 nm. One-sixth of the volume of an embedded triangle corresponding to the monomer volume was calculated to be 18 $nm^3$, and the mass in this box was assumed to be 9963 Da from the protein monomer. The shell density is taken to be the density of the monomer.

Impact of variations in shell parameter choices on inferred mass: As we see in the main text, the total estimated mass of a carboxysome depends primarily on the scattering cross-section, and secondarily on the total radius. As we show below, the distribution of mass within this radius (partitioning between core and shell) does not affect the total mass estimation significantly. This can be seen in the fact that the estimated masses for the carboxysomes studied here do not significantly vary with small variations in the shell thickness and density in the range 2.75 nm → 3.25 nm → 3.75 nm, and 0.70 g/mL → 0.92 g/mL → 1.14 g/mL respectively (See Fig S8). The major effect of increasing shell thickness and density is to create a larger range of unphysical carboxysomes with large radius and small scattering cross-section, since with increasing shell mass, there is a decrease in the minimum scattering cross-section for any given radius.

**Note S5: Correlated distributions of carboxysome scattering cross-section and radius**

Generating correlated distributions: The analysis of the core mass *vs* normalized scatter trend requires some method to parametrically vary between fully uncorrelated to strongly correlated carboxysome scattering cross-section and radius values. Put another way, we need to generate distributions of carboxysome parameters where the scattering cross-section and the radius distributions match the measured distributions, including the effect of a trend between the two. We use Gaussian copulas to generate these correlated samples.[1] Briefly, we generate a pair of

Gaussian random variables with a known correlation function. We then transform the random variables to have marginal distributions equal to the measured distributions of our two desired variables, $\sigma_{scat}$ and r. Since the case of correlation = 0 and correlation = 1 between the Gaussian variables map onto our desired extreme cases, we can use intermediate values of correlation to move between the two extremes. This scheme is outlined in Figure S9.

The initial Gaussian variables $(Z_1, Z_2)$ are generated with a correlation $R$ enforced by setting the covariance (0.6 for Figure S9). Applying the cumulative density function (CDF) $CDF_{\mathcal{N}(0,1)}$ of the standard Gaussian distribution to each of these variables gives two uniform variables $(U_1, U_2)$ that still have some underlying correlation. We then want to relate these uniform variables to the experimental distributions for $r$ and $\sigma_{scat}$. The transformation from a random number $U$ on the interval $[0, 1]$ to any variable $X$ is given by the inverse CDF of the variable, $X = CDF_X^{-1}(U)$. This inverse CDF can be easily estimated for a measured distribution. Each sampled value $X$ in the sorted (low to high) distribution is associated with its normalized index $U$ in [0, 1]. For example, for samples $x_i, i = 1 \dots N$ of a random variable $X$ sorted such that $i < j \Rightarrow x_i \leq x_j$, we estimate $CDF_X^{-1}\left(\frac{i}{N}\right) = x_i$. These discrete values are interpolated to produce a continuous function. We estimate $CDF_{\sigma_{scat}}^{-1}(u)$ and $CDF_r^{-1}(u)$ from our measured distributions in Figure 5d and Figure 5e of the main text. These functions can then be applied to $U_1$ and $U_2$ respectively to obtain our random samples of $\sigma_{scat}$ and $r$ including the assumed underlying correlation. For an initial correlation of R = 0.6 of the two Gaussian variables, we compute a Pearson's correlation coefficient of 0.561 between $r$ and $\sigma_{scat}$. We refer to these correlated distributions by the covariance, R, used to generate the Gaussians.

<u>Plotting density map of the correlated distribution:</u> The coloring of points in Figure 6a,d,g represent the data density calculated as the reciprocal sum of squared distances from each sampled point on the matrix of values.
